# The structure of full-length human phenylalanine hydroxylase in complex with tetrahydrobiopterin

**DOI:** 10.1101/552281

**Authors:** Marte Innselset Flydal, Martín Alcorlo-Pagés, Fredrik Gullaksen Johannessen, Siseth Martínez-Caballero, Lars Skjærven, Rafael Fernandez-Leiro, Aurora Martinez, Juan A. Hermoso

## Abstract

Phenylalanine hydroxylase (PAH) is a key enzyme in the catabolism of phenylalanine, and mutations in this enzyme cause phenylketonuria (PKU), a genetic disorder that leads to brain damage and mental retardation if untreated. Some patients benefit from supplementation with a synthetic formulation of the cofactor tetrahydrobiopterin (BH_4_) that partly acts as a pharmacological chaperone. Here we present the first structures of full-length human PAH (hPAH) both unbound and complexed with BH_4_ in the pre-catalytic state. Crystal structures, solved at 3.18 Å resolution, show the interactions between the cofactor and PAH, explaining the negative regulation exerted by BH_4_. BH_4_ forms several H-bonds with the N-terminal autoregulatory tail but is far from the catalytic Fe^II^. Upon BH_4_ binding a polar and salt-bridge interaction network links the three PAH domains, explaining the stability conferred by BH_4_. Importantly, BH_4_ binding modulates the interaction between subunits, providing information about PAH allostery. Moreover, we also show that the cryo-EM structure of hPAH in absence of BH_4_ reveals a highly dynamic conformation for the tetramers. Structural analyses of the hPAH:BH_4_ subunits revealed that the substrate-induced movement of Tyr138 into the active site could be coupled to the displacement of BH_4_ from the pre-catalytic towards the active conformation, a molecular mechanism that was supported by site-directed mutagenesis and targeted MD simulations. Finally, comparison of the rat and human PAH structures show that hPAH is more dynamic, which is related to amino acid substitutions that enhance the flexibility of hPAH and may increase the susceptibility to PKU-associated mutations.

**Significance Statement:** The present crystal structure of phenylalanine hydroxylase (PAH) provides the first and long-awaited 3D-structure of the full-length human PAH, both unbound and complexed with the tetrahydrobiopterin (BH_4_) cofactor.

The BH_4_-bound state is physiologically relevant, keeping PAH stable and in a pre-catalytic state at low L-Phe concentration. Furthermore, a synthetic form of BH_4_ (Kuvan^®^) is the only drug-based therapy for a subset of phenylketonuria patients.

We found two tetramer conformations in the same crystal, depending on the active site occupation by BH_4_, which aid to understand the stabilization by BH_4_ and the allosteric mechanisms in PAH. The structure also reveals the increased mobility of human-compared with rat PAH, in line with an increased predisposition to disease-associated mutations in human.

## Introduction

Phenylalanine hydroxylase (PAH) catalyzes the tetrahydrobiopterin (BH_4_)-dependent hydroxylation of L-phenylalanine (L-Phe) to L-tyrosine (L-Tyr), which is the rate-limiting step in the degradation of L-Phe and L-Tyr. In humans this reaction is vital for preventing the accumulation of neurotoxic amounts of L-Phe, which is the hallmark of the disease phenylketonuria (PKU) (1-3). Patients must follow a life-long protein-controlled diet to avoid severe neurological symptoms, but in a subset of the patients the diet can be alleviated by sapropterin dihydrochloride (Kuvan^®^), a synthetic form of the cofactor BH_4_ (1) that shows multifactorial effects on PAH mutants, including their stabilization (4). As most of the mutations in *PAH* cause protein instability, leading to misfolding and loss-of-function (3), it is of medical interest to discover and optimize other compounds with pharmacological chaperone potential that can specifically prevent PAH misfolding and extend the range of mutations that can be rescued (5, 6). Structural information is essential in this endeavor.

Mammalian PAH is tetrameric and its 52 kDa subunits are composed of a regulatory domain (RD, residues 1-110) with an unstructured N-terminal tail (N-term; residues 1-29), a catalytic domain (CD, residues 111-410), and an oligomerization domain (OD, residues 411-452) responsible for dimerization and the subsequent tetramerization (SI Appendix, Fig. S1). This oligomeric and multidomain arrangement allows the complex regulation of PAH activity, including *(i)* activation by the substrate L-Phe, which elicits a positive cooperative response on PAH ensuring that neurotoxic accumulation of L-Phe is avoided, and *(ii)* decreasing PAH activity at low substrate concentration to preserve a lower threshold value of L-Phe for protein synthesis (for reviews see (2, 7, 8)). At low, physiological concentration of L-Phe, PAH is complexed with the cofactor BH_4_ (present in hepatocytes at equal concentrations as PAH subunits) that acts as a negative regulator of the activation by L-Phe (8, 9). BH_4_ also stabilizes PAH and increases the steady-state levels of the enzyme *in vivo* (10, 11). While the activation by L-Phe induces a large conformational change (8), recently proposed to involve dimerization of RDs (12-14), BH_4_ binding elicits a limited conformational change proposed to be mostly constrained to the N-term tail (15, 16). The importance of these N-terminal 29 residues in both the inhibitory regulation by BH_4_ and the activation by L-Phe is evident, as PAH lacking this tail is not regulated by either BH_4_ or L-Phe and is constitutively active (15).

Until now, the available mammalian PAH structures are truncated, lacking one or two of the domains (17-19) or full-length without ligands, the latter only of rat PAH (rPAH) (12, 14). Understanding of the structural defects associated to PKU mutations and development of specific stabilizing therapies would highly benefit from the availability of the structure of hPAH, both with and without ligands. Thus, we aimed to solve the long-awaited full-length structures of hPAH both in the absence of ligands and with the cofactor BH_4_ and have obtained two structures at 3.18 Å resolution from the same crystal, which present different occupancy by BH_4_. The full-length structure in absence of BH_4_ has been also obtained at medium resolution by cryoEM. These are the first structures of complete hPAH with all domains and the first structures of full-length mammalian PAH with ligand.

## Results and discussion

### Crystallization of hPAH with two tetramers in a single crystal

Both full-length hPAH and ΔN13-hPAH, the latter produced by cutting MBP-hPAH with Factor Xa protease (20, 21), were co-crystallized with BH_4_ and produced crystals with similar appearance at the same conditions (see *Materials and Methods*). However, data sets collected from hundreds of similar looking crystals showed both low resolution and severe anisotropy, suggesting that there was still heterogeneity in the conformation of tetrameric hPAH. Nevertheless, with ΔN13-hPAH and DTT we eventually obtained a crystal with a degree of anisotropy that allowed us to obtain the structural determination of hPAH at 3.18 Å resolution (Fig. 1, SI Appendix, Fig. S2 and Table S1) (22). Comparison with the structure of the dimeric catalytic domain (hPAH-CD) – that we also solved at 1.67 Å resolution (SI Appendix, Table S1) with previously published conditions (17) (SI Appendix, Fig. S2B) – shows that the absence of Fe in our structure does not alter the arrangement of the iron-binding 2-His-1-carboxylate facial triad (Fig. 1C). Four independent monomers of PAH were found in the asymmetric unit belonging to two different tetramers, Tet1 and Tet2, each created as a dimer of dimers by a two-fold crystallographic axis. While Tet1 presents the BH_4_ bound in two active sites (chains A, B) and the other two (chains C, D) without the cofactor, Tet2 shows BH_4_ bound to all four active sites (Fig. 1). All four independent monomers in both tetramers present a similar overall structure for the backbone (RMSD ∼0.6 Å for Cα superimposition), with hPAH arranged as a tetramer through the C-terminal oligomerization helix, following an approximate 222 symmetry between monomers, only the two-fold symmetry between dimers being strict (Fig. 1A). Each monomer presented electron density for all domains (residues 22-450) (Fig. 1A, SI Appendix, Fig. S1). The tetramers can be roughly encased in a rectangle with a short edge along the dimer (∼70 Å) and a long edge (∼110 Å) between the two dimers, while differences in the long edges are observed between Tet1 and Tet2 (SI Appendix, Fig. S3). The RD is essential for this specific oligomeric arrangement in hPAH as deduced by comparison with the human truncated form lacking RD (PDB 2PAH) (18) (SI Appendix, Fig. S4).

**Fig. 1.**
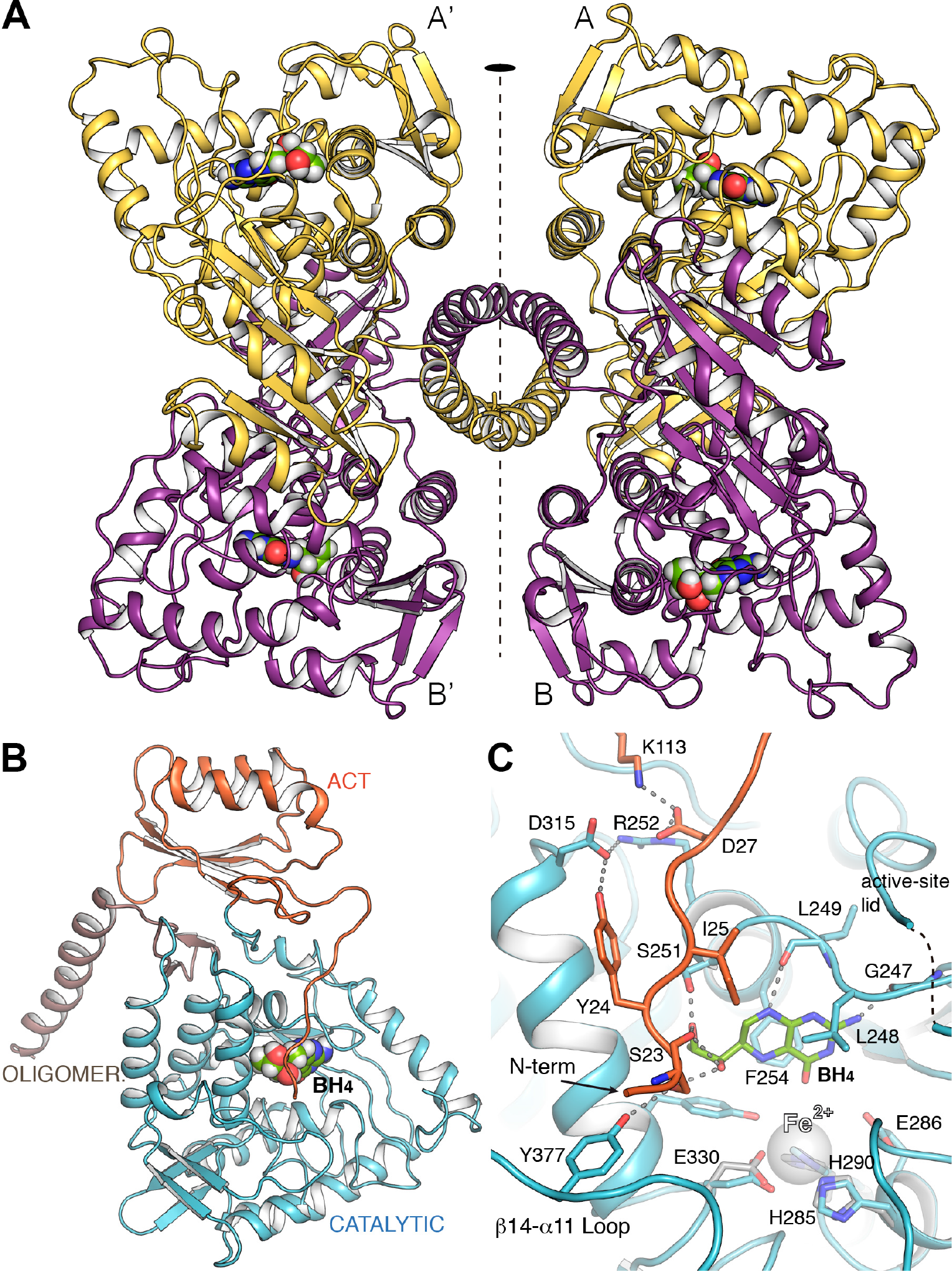
Crystal structure of human PAH. (**A**) Ribbon representation of hPAH in complex with BH_4_ at the four active sites (Tet2). The tetramer is formed as a dimer of dimers by a two-fold symmetry (depicted). BH_4_ cofactor is drawn as spheres. (**B**) The hPAH monomer with each domain colored differently. (**C**) Detailed view of the active site of hPAH in complex with BH_4_ (green sticks). Relevant residues are represented as capped sticks and labeled. The Fe^II^ cation, not present in the structure, is represented as a transparent sphere for comparison purposes. Its location is obtained by direct superposition with the CD of human PAH (obtained in this work). Residues from this structure involved in the interaction with Fe^II^ cation are shown in gray sticks.

The presence of unbound and BH_4_-bound states – from here on referred to as apo and holo states, respectively – in the same crystal allowed a direct comparison between both conformations. Until now, investigations of how cofactor binding affects the conformation of the RD have come from modelling through superimposition of truncated structures (apo rPAH containing RD and CD, PDB 1PHZ) onto hPAH-CD with ligands (i.e. with BH_4_, PDB 1J8U, and with both BH_4_ and with substrate analog thienylalanine, THA, PDB 1MMK). These modelling analyses have predicted interactions of BH_4_ with S23 at the flexible and unstructured N-term (15, 16, 23), a segment that has been termed an intrasteric autoregulatory sequence (IARS) (19). Comparing our apo and holo structures with each other and with the previous ligand-bound truncated forms reveals main events implicated in the negative regulation of PAH by BH_4_ and derived enzyme stabilization, as well as in the conversion to the active BH_4_-bound structure in the presence of substrate. As detailed below, the conformational events involve so far unidentified networks of residues, far beyond the IARS.

### Structural effects of BH_4_ binding associated to negative regulation of activation by L-Phe and enzyme stabilization

The BH_4_ binding-site is flanked by the N-term (residues 21-32), the active-site lid (130-150), the Fe^II^-coordinating residues, the β6-α7 loop (residues 245-251) and F254 (Fig. 1C). BH_4_ is sandwiched between hydrophobic residues, F254 from one side and L248 and I25 from the other (Fig. 1C). Besides, some polar interactions are observed between BH_4_ and residues from the IARS and the β6-α7 loop (Fig. 1C). The direct interaction of S23 in the N-term with BH_4_, via H-bonds with both O1’ and O2’ in the dihydroxypropyl side chain, is observed for the first time in our BH_4_-bound structure, and confirms previously predicted interactions with these hydroxyls in the natural (6*R*)-BH_4_ configuration (15, 16, 23). These interactions appear related to the inhibitory effect of BH_4_ on L-Phe activation of PAH, and explain why the cofactor analogue 6-methyl-tetrahydropterin (6-MPH_4_) is not inhibitory and (6*S*)-BH_4_ is less inhibitory (24).

The activation of PAH by L-Phe requires a slow (s-to-min timescale with 1 mM L-Phe, depending on the temperature) global conformational change, and the activation rate is slower for the BH_4_-preincubated than for the unbound enzyme (8-10, 23), associated to the negative regulation of PAH activation exerted by BH_4_. Recent structural analyses have shown that full activation involves the shift and dimerization of the RDs, with concomitant movement of the N-term IARS that releases from the CD (12, 14). In apo hPAH, S23 H-bonds with Y377 in the CD, whereas in the holo structure S23 is also engaged in a larger H-bond network involving BH_4_ and extends to residues in the CD and the RD (Fig. 1C). Structural comparison between apo- and holo-hPAH monomers reveals other BH_4_-induced changes, beyond the N-term IARS (Fig. 2A). Surprisingly, the conformation of the N-term is similar for apo and holo hPAH (Fig. 2A), and also for apo rPAH (12), showing that binding of BH_4_ does not induce large conformational changes on the N-term. Comparative analyses of apo- and holo-hPAH monomers by calculating differences in inter-residue distances between the apo and all holo monomers (see *Materials and Methods*) revealed many distances that facilitate or abolish interactions upon BH_4_ binding, most involving residues in the CD (SI Appendix, Fig. S5 and Table S2). Out of the 35 largest distance differences, about half engage intra-CD residue pairs (19 pairs), most concerning residues around the BH_4_ binding site (SI Appendix, Table S2). These are followed by intra-RD residues (6 pairs), between RD and CD (5 pairs), between RD and OD (3 pairs), and intra-OD and between CD and OD (1 pair each). Thus, upon BH_4_ binding changes are observed that link the CD with the OD, both directly and via the RD. Actually, the largest change (∼6 Å) is the increased distance between K73 in the RD and L430 at the start of the tetramerization helix in the OD, indicating conformational changes affecting oligomerization and allostery (see below). In the active site, the most apparent change involves residues L248 and F254 in the BH_4_ binding-site (Fig. 2A1 and SI Appendix, Table S2). In the apo subunit, these two residues show a direct van der Waals interaction (∼3 Å), observed for the first time, and partly occupy the BH_4_ binding-site, whereas in the BH_4_-bound active site they move to sandwich the cofactor (Fig. 2A). Interestingly, binding of BH_4_ changes the interaction network between the N-term and the CD (Fig. 2A1). In the apo state residue D27 makes H-bond interactions with α7 residues (notably S251) while in the holo state makes a salt-bridge interaction with R252 and K113 instead. The salt-bridge interaction between E26 and K113 observed in apo state is lost in holo state (Fig. 2A1, SI Appendix, Fig. S5). Interactions of BH_4_ with the N-term and the CD, as well as the network of polar and salt-bridge interactions linking the RD (Y24, D27, K113) with the CD (D315, R252) (Fig. 1C), appear determinant for the regulatory inhibitory effect and the stabilization of BH_4_. Differential scanning calorimetry (DSC) experiments at 200 °C/h with the BH_4_ analog 7,8-dihydrobiopterin (BH2) (SI Appendix, Fig. S6A) support the stabilization effect of BH_4_ beyond the CD. Hence, while apo-hPAH presents the expected unfolding transition temperatures at ∼49 and 58 °C, previously associated to the RD and the CD, respectively (27), both transition temperatures increased ∼2 °C in holo-hPAH (SI Appendix, Fig. S6A).

**Fig. 2.**
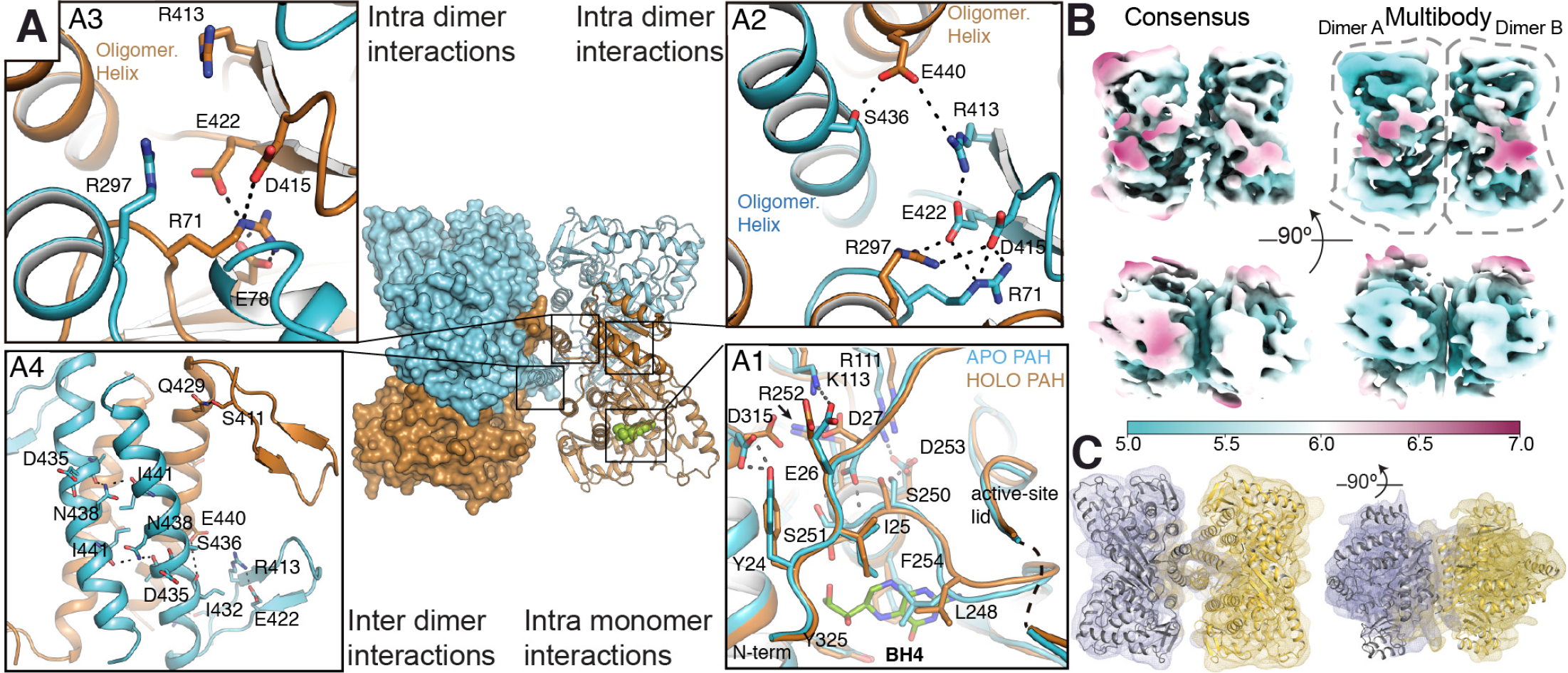
Long-range effects in hPAH upon BH_4_ binding and cryo-EM of apo-hPAH. (**A**) Tet1 with monomers colored differently (apo in cyan and holo in ochre). (**A1**) Structural comparison of the active site in apo hPAH *vs* BH_4_-bound hPAH. (**A2**,**A3**) Detailed view of the apo-holo dimer interface showing network of salt-bridge interactions. (**B**) Top and side view of the cryo-EM reconstruction of apo-hPAH tetramer (left) and multi-body reconstruction of two independent dimers (right). Map color represents the map local-resolution estimation (see color guide). Note that the multi-body refined map is a composite of two dimers refined independently. (**C**) Rigid body structural fitting of Tet2 dimer onto the cryo-EM multi-body reconstruction of two independent dimers.

### Long-range structural effects of BH_4_ binding

A different disposition of the monomers in Tet1 and Tet2 is observed depending on occupancy by BH_4_ that changes the overall dimensions of the tetramer, the distances between monomers and the relative disposition (∼5 Å gliding) of one dimer versus the other through the oligomerization helices (SI Appendix, Fig. S3). Considering that all monomers presented a nearly identical backbone we reasoned that the observed differences should rely on different structural arrangements inside the dimers that would promote changes in the tetramer. Superposition of the holo monomers in apo-holo and holo-holo dimers shows a rotation of ∼3° and displacements up to 3 Å between monomers. Thus, the changes observed inside the active site of a monomer upon BH_4_ binding (Fig. 2A1) must also propagate to the dimeric partner (intra dimer interactions) and to the tetramer (inter dimer interactions). Structural analysis of Tet1 and Tet2 provides clues about such long-range effects of BH_4_ binding (Fig. 2A). Changes are observed at the dimer interface between the apo-holo or holo-holo dimers (SI Appendix, Fig. S7). It is worth to mention residue R297 that participates in an intra-dimer interaction network with residues E422, D415 (OD) and R71 (RD) only when BH_4_ is bound (Fig. 2A2, 2A3, SI Appendix, Fig. S7). Another relevant residue in this network is R413 that, in the apo monomer, can establish a salt-bridge interaction with the oligomerization helix (residue E440) of the dimeric partner (Fig. 2A2). This interaction is not seen between holo monomers (SI Appendix, Fig. S7). As previously mentioned, the quaternary structure of hPAH is also affected by BH_4_ binding, the most significative is a gliding of ∼5 Å between dimers along the OD helices (Sup. movies 1 and 2) changing the network of polar interactions between the OD helices in the tetramer (Fig. 2A4, SI Appendix, Fig. S8). Contrary to tyrosine hydroxylases or tryptophan hydroxylases the OD helices in mammalian PAH lack a proper Leu zipper and present abundance of polar residues inside the coiled-coil structure, as has already been noticed in the structure of rPAH (12). The lack of a classic leucine zipper explains why PAH can dissociate to dimers as well as the conformational variability in the C-terminal helices.

To further investigate the dynamics of apo hPAH tetramer, i.e. with all four subunits in the apo state, we performed single particle cryo-electron microscopy (cryo-EM) studies on the same full-length construct but in the absence of cofactor (see *Materials and methods* and SI appendix Table S3). A consensus refinement of the hPAH tetramer yielded a cryo-EM map of resolution ranging from 5 to 7 Å due to the high mobility of the monomers within the tetramer (Fig. 2B). 3D classification allowed for the separation of different conformations in the dataset (SI Appendix, Fig. S9), but these still harbored the same flexibility-related problems as the consensus refinement. In order to address these flexibility issues, we performed multibody refinement by masking individual dimers (Fig. 2B) (29). This procedure improved resolution to ∼5 Å for most of the protein except for the RD (Fig. 2B) resulting in an excellent fit of the crystal structure onto the resulting maps (Fig. 2C). Moreover, variance analysis using principal component analysis showed that most of the orientational differences between dimers can be described by three main components (SI Appendix, Fig. S10). Interestingly, the amplitude distribution along these components is monomodal, indicating that these are continuous movements in which dimers rotate around each other mainly as rigid bodies (SI Appendix, Fig. S10C and Sup. movie 3). Thus, in absence of BH_4_, hPAH dimers present a large range of motion inside the tetramer. While there are no noticeable changes in the internal structure of the CD, the RD seems to present extra flexibility.

In agreement with crystallographic and cryo-EM data, MD simulations (eight 100-ns long runs) of hPAH performed with and without BH_4_ (see *Materials and methods* and SI Appendix, Fig. S11) show a highly dynamic tetrameric structure in both apo and holo states, with large-scale inter-dimer motions, but relatively small intra-dimer fluctuations. We calculated the fluctuation profile (root mean square fluctuation; RMSF) for each monomer in each of the simulations. Comparative analysis shows that BH_4_ stabilizes significantly (*p < 0.05*) several regions in the CD, but increases flexibility in loop α14-β11 (residues 374-384) that contains Y377 residue (Fig. 1C, SI Appendix, Fig. S12). This loop, important for embracing the substrate, moves ∼5 Å closer to the active site upon substrate binding (30). The Y138 loop, termed the active site lid, also moves considerably (> 10 Å) towards the active site upon L-Phe binding (30) but, contrary to loop α14-β11, it loses mobility upon BH_4_ binding (SI Appendix, Fig. S12).

### The significance of the PAH:BH_4_ pre-catalytic complex and the role of Y138

In the holo hPAH subunits, BH_4_ is bound placing its O4 atom, the site of hydroxylation of the pteridine ring during catalysis, at 3.8 Å from the catalytic iron (Fig. 3A), 0.7 Å longer than the required distance to form the Fe^II^-OO-BH_4_ adduct and generation of the Fe^IV^=O hydroxylating intermediate (25, 30, 31). The cofactor bound hPAH structure presented in this work thus corresponds to a pre-catalytic conformation, which is of high significant physiological relevance, as PAH and BH_4_ coexist in hepatocytes in stoichiometric amounts, forming a stable complex at normal low concentrations of L-Phe *in vivo* (≤ 54 µM) (10). BH_4_-bound hPAH presents lower initial catalytic activity, and is activated by L-Phe at a slower rate compared with the ligand-free enzyme (8-10, 23) (SI Appendix, Fig. S6B and see below). The physiological significance of the negative regulation of PAH activation by BH_4_ is related to the maintenance of a low hydroxylating activity at low substrate concentrations to preserve a basal level of L-Phe required for protein synthesis (9, 10). Furthermore, the stabilization of PAH by BH_4_ binding (SI Appendix, Fig. S6A) contributes to maintaining the *in vivo* half-life of the enzyme, as supported by the decrease of steady state levels of PAH caused by BH_4_ loss in cells and in liver (11, 32, 33). No atomic structural information is available on the large conformational changes associated to complete L-Phe activation, including the dimerization of the ACT domains (12, 14). Nevertheless, our structures provide novel insights into the substrate-induced conformational changes around the active site related to the transition from the pre-catalytic to the catalytic state upon entrance of the substrate at the active site. Superposition of holo-hPAH with the binary hPAH-CD:BH_4_ and ternary hPAH-CD:BH_4_:THA complexes reveals a possible role of Y138 in this transition by interacting with the O1’ hydroxyl and contributing to the displacement of the cofactor 0.7 Å and 2.6 Å towards the iron and Glu286, respectively (Fig. 3A,B). In order to probe plausible transition pathways occurring upon substrate binding, we performed a 2 ns long targeted MD (TMD) simulation of Tet2 in complex with L-Phe and BH_4_ (Sup. movie 4). With this simulation technique we gradually steer one catalytic domain of Tet2 to a target structure defined by the ternary hPAH-CD:BH_4_:THA complex (PDB 1MMK). Of note, the binding site of L-Phe was solvent accessible and there was no clash of the bound substrate with residues in hPAH. At ∼0.8 ns of the simulation Y138 has moved ∼10 Å from its original position towards the active site and forms a transient H-bond with the carbonyl O of P279. After ∼1.4 ns (Fig. 3C), a H-bond is formed between the side chain of T278 and the amino group of L-Phe, and the translation of Y138 towards the active site is accelerated, and at the end of the simulation, a H-bond is formed between the phenol O of Y138 and the O1’ of BH_4_ in its pre-catalytic position (Fig. 3D). At that point the cofactor only maintains a single H-bond interaction with S23, which gets further weakened at the end of the simulation when S23 interacts with S251. To further investigate the role of the hydroxyl-group of Y138 in initiating the movement of BH_4_ towards the catalytic binding mode, as well as the relevance of S23 and its interacting CD-residue Y377 (Figs. 1C, 3A-D), we prepared the hPAH mutants Y138F, S23A and Y377F, which were expressed and purified as tetramers, but at lower yield than wild-type (WT). Other residues that interact with BH_4_ in the pre-catalytic conformation (region 245-255, 286, 322 and 325), also interact with BH_4_ in the catalytic conformation and, in addition, these sites are actually associated with severe destabilization of PAH (www.biopku.org), and thus not suitable for mutagenesis investigations. The initial reaction rates at 37 °C, comparative to the hPAH-WT, were measured for the four enzymes in 3 initial states, i.e. *L-Phe*-*activated* (preincubated with 1 mM L-Phe), *non-activated* (not preincubated with either substrate or cofactor) and *pre-catalytic BH_4_-bound* (preincubated with 75 µM BH_4_) (Figs. 3E, SI Appendix, Fig. S6B). The concentration of substrate and cofactor were otherwise identical at the start of the reactions. Activated Y138F shows reduced activity, to 70% of WT, but more remarkable is the large reduction in activity for the non-activated and pre-catalytic states for this mutant (to 35-45% of WT). This reduction is not associated to different steady state kinetic parameters, as the Y138F mutant presents similar *K*_m_(BH_4_), *S*_0.5_(L-Phe) and hill coefficient (*h*) as hPAH-WT (SI Appendix, Fig. S6C), supporting a role of the Y138 hydroxyl group in disengaging BH_4_ from the pre-catalytic conformation in the full-length enzyme. The other two mutants S23A and Y377F were prepared to validate the effect of the H-bonding interactions centered at S23 to stabilize the enzyme-cofactor complex in the pre-catalytic state. Both mutants, and notably S23A, show reduced activity for the L-Phe-activated state and increased *S*0.5(L-Phe) (SI Appendix, Fig. S6C), indicating the participation of these residues in substrate activation, through the side-chain O in the case of S23. Moreover, the mutant S23A is expected to maintain the negative regulatory interactions with O1’ and O2’ of BH_4_ through its backbone carbonyl and amide groups, explaining the low basal activity of the BH_4_-bound state. On the other hand, as the function of Y377 appears related to its interaction with S23 for proper localization of this residue and the IARS through H-binding interactions (Fig. 3A-C), the rupture of the H-bond with S23 in the Y377F mutant results in an increased activity for the unactivated states (Fig. 3E, SI Appendix, Fig. S6B).

**Fig. 3.**
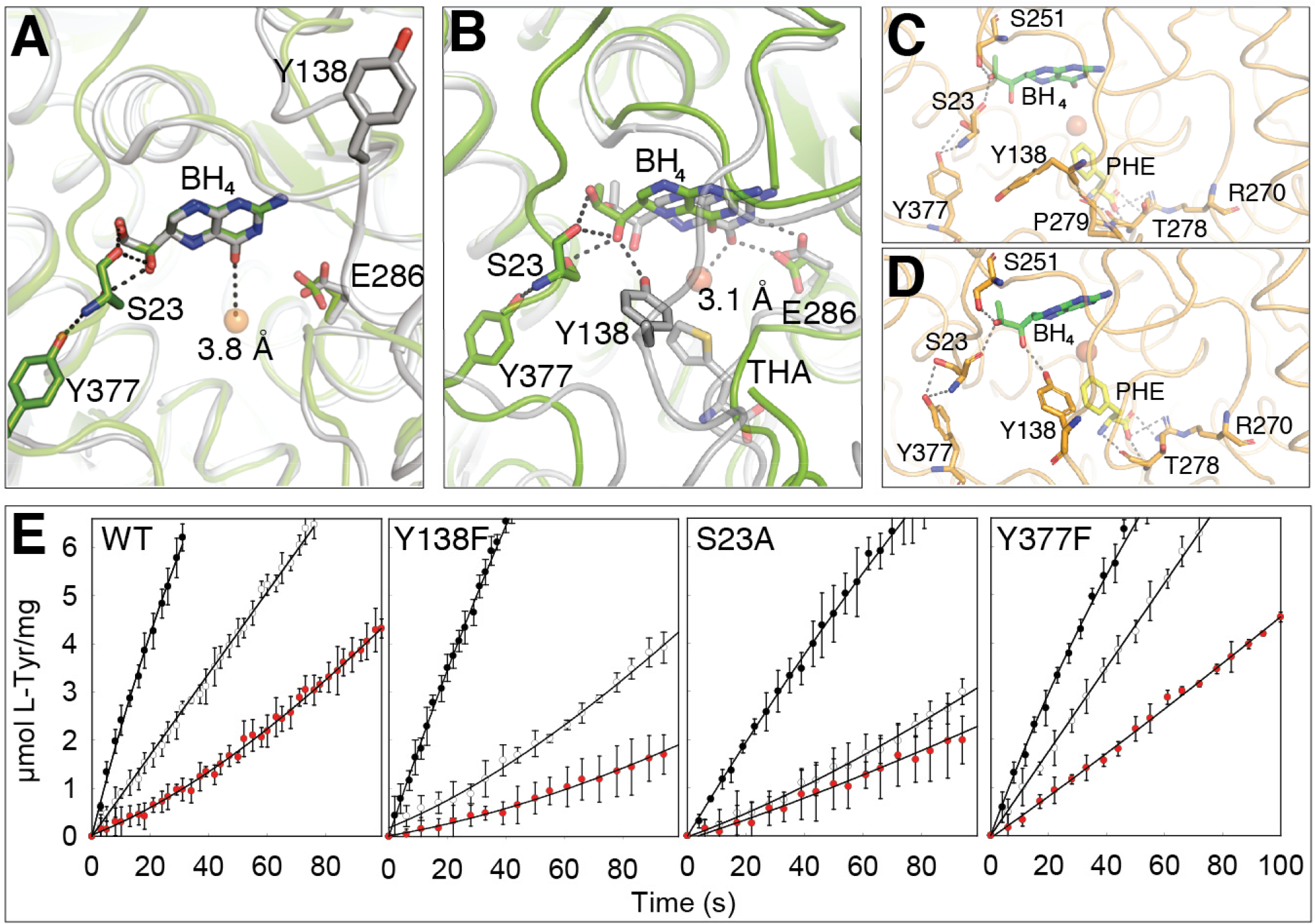
Transition to active cofactor binding. (**A**) Superposition of hPAH:BH_4_ (Tet2A; green) with hPAH-CD:BH_4_ (PDB 1J8U; grey). In both, BH_4_ is in pre-catalytic position, with the O4 atom 3.8 Å from the Fe^II^ and Y138 (not visible in Tet2) ∼20 Å away. H-bonds shown as dashed lines. (**B**) Superposition of hPAH:BH_4_ (Tet2A; green) with hPAH-CD:BH_4_:THA (PDB 1MMK; grey). BH_4_ is at the catalytic position, interacting with E286 and with the O4 3.1 Å from Fe^II^ and generates the Fe^II^-O2-BH_4_ adduct. The movement of Y138, induced by substrate analogue (THA) binding, positions the hydroxyl group of Y138 2.9 Å away from O1’ in the dihydroxypropyl side chain of BH_4_. (**C,D**) Snapshots from the TMD of L-Phe binding to full-length hPAH (Tet2A). At ∼1.41 ns, L-Phe H-bonds to T278, and the translation of Y138 is accelerated (C). At the end of the 2 ns-simulation the phenol O of Y138 H-bonds with the O1’ of BH_4_ (D). (**E**) Initial reaction rates for hPAH-WT and mutants, starting the reaction as *L-Phe*-*activated* (black circles), *non-activated* (white circles) and *pre-catalytic BH_4_-bound* (red circles).

### Human and rat PAH present significant structural differences despite high sequence homology

PAH for human and rat share a high sequence similarity (96%). Among the 452 residues of hPAH there are only 16 non-conservative- and 14 conservative mutations compared with rPAH (SI Appendix, Fig. S13). Conservative mutations are distributed among the three domains and non-conservative mutations are located in RD and CD. Considering differences in size of RD and CD, mutations are strongly concentrated in the RD (8% of residues of RD are mutated between hPAH and rPAH *vs.* 2% of the CD) supporting differences in the regulatory properties of both enzymes (34).

Overall structures of the human and rat PAH monomers are very similar (RMSD values for the backbone ranging 0.8-1.1 Å) (SI Appendix, Fig. S14A). Major differences are seen at the interface between RD and CD (loop β1-α1 (44-46)), in α2 (88-100) of the RD, residues 115 in the hinge region 110-118 (loop β4-β5), 143, 338 in the CD, and at the beginning of the OD helix (L427-K438) (SI Appendix, Fig. S14, S15) that provoke differences in tetramer arrangement (SI Appendix, Fig. S15). Regions with largest conformational changes in the RD are located around non-conservative mutations, i.e. C29 (S29 in rPAH), L88 (K88 in rPAH), K114 (E114 in rPAH), and D116 (N116 in rPAH) (SI Appendix, Fig. S14B). In the RD/CD interface is worth to mention the salt-bridge in rPAH between E114 and K88 (K114 and L88 in hPAH), linking α2 with loop 110-118, that obviously is absent in hPAH, where a different interaction pattern is observed in this region. In the apo-state of hPAH salt-bridges are observed between D122 and both K114 and K85 and D116 and R99. Consequently, both sides of the α2 helix in the RD, i.e. the C-terminal (K99) side in hPAH and the N-terminal (K85/K88) side in rPAH, are involved in interactions with the important loop 110-118, located at the end of the RD, at the boundary between the RD and the CD. Differences are also observed in the conformation of the OD helices, i.e. in rPAH the C-terminal helices of the four monomers adopt two distinct orientations with respect to the CD, differing by a tilt of ∼10°, and with the monomers situated across the diagonal of the tetramer containing similarly positioned helices (12). Structural analysis reveals that the different conformations relate with a different pattern in polar interactions at the beginning of the OD helices depending on whether Q428 is making a H-bond with N73 or inserted in the interface between the CD and the RD (SI Appendix, Fig. S15). In hPAH there are no such distinct orientations for the C-terminal OD helices that are placed in between the two conformations observed in rPAH (SI Appendix, Fig. S14). In hPAH the interaction network in this region is affected by the presence of the BH_4_ cofactor in the active site. In the apo Q428 makes H-bond interactions with both the backbone atoms of the region connecting the OD helix with the protein core and with residue E78 (SI Appendix, Fig. S14C), while in the holo Q428 interacts with S70 and, interestingly, K73 (N73 in rat) is salt-bridging with D425, thus contributing to the locked, inhibited structure in the holo-state of hPAH.

The effect of mutations between human and rat PAH was studied by parallel MDs with our hPAH structures and existing rPAH structure(s) (see *Materials and methods* and SI Appendix, Fig. S16). In agreement with our structural results, fluctuation analyses showed that, overall, hPAH is more dynamic, notably in the regions with significant structural differences (SI Appendix, Fig. S17), such as RD, showing non-conservative mutations (SI Appendix, Fig. S13B), and the area around the OD helix.

### Relevance for HPA and BH_4_-responsive PKU

Increased understanding of the high vulnerability of human enzymes - notably those highly regulated, associated with central metabolic pathways - to disease-associated mutations points to evolutionary divergence from consensus amino acids that confer stabilization (35). This evolutionary mechanism also appears to describe the adaptation of hPAH towards a less stable and highly mobile enzyme (SI Appendix, Fig. S17), most probably associated with the acquisition of the flexibility required for prompter response to changes in substrate and cofactor concentration along the evolution of the neuronal system (2). Flexibility is however often traded off with an increased susceptibility to mutations (35). Indeed, for the more than 900 hPAH mutations causing HPA/PKU (www.biopku.org), several studies have shown the relationship between hPAH protein stability and both remaining activity and allelic phenotype (3, 36). The enzyme destabilization caused by each mutation is actually highly predictive of patient phenotype and therapeutic response to BH_4_ supplementation (3). As here reported, the stabilization by BH_4_ is initiated at its binding-site in the CD but also affects the whole structure through networks that propagate to the RD and OD both inside the monomer (Fig. 2A, SI Appendix, Fig. S5) and intra dimer (Fig. 2). HPA/PKU-associated mutations are located throughout the hPAH structure, but seem to be concentrated in hotspot regions with highly destabilizing mutations (235–281, corresponding to exon 7, and 282–330 to exons 8 and 9 in the *PAH* gene) (36). These regions include the flexible loop β6-α7 (244-250), whose conformation is modified (Fig. 2A1 and SI Appendix, Fig. S5) and rigidified (SI Appendix, Fig. S12) upon BH_4_ binding and presents a high mutation frequency with several mutations associated with moderate-to-severe (residues P244, V245, A246, G247 and L249) or mild (L248) HPA/PKU phenotype (www.biopku.org and (36)).

The hotspot regions also include mutations at R252, F254, and D315 (involved in changes from apo to holo conformation) (www.biopku.org and (36)). Together with S251, R252 appears essential for the stabilizing networks established with RD residues D27 in the IARS upon BH_4_ binding (Fig. 2A1, SI Appendix, Fig. S5). Likewise, the propagation of BH_4_ binding from the CD to the OD through the RD towards the other subunits in the dimer also involves regions with high mutation frequency, such as the β2-β3 loop, which interacts with R297, D415 and E422 (Fig. 2A).

In conclusion, the identification of the networks that transmit the BH_4_-bound state from the binding site at the CD to the other regions in the monomer, dimer and tetramer of hPAH, increments our understanding of the negative regulation of PAH by its cofactor and the stabilization caused by BH_4_-binding, observed both for WT and mutant hPAH (4, 11). Thus, the structures of hPAH tetramers totally and partially bound with BH_4_ presented here provide a rationale for BH_4_-responsive PKU by sapropterin dihydrochloride, and pave the way to new stabilizing/chaperoning therapeutic approaches to address PKU.

## Materials and Methods

Descriptions of the protein production; crystallization, crystallographic and cryoEM structural determinations; of the computational methods; of activity measurements, and differential scanning calorimetry; and of the reagent preparations are in SI Appendix.

## Accession numbers

The atomic coordinates for hPAH have been deposited in the PDB with codes 6HYC and 6HPO, for the full-length hPAH and catalytic domain respectively. The EM maps have been deposited in the EMDB with accession s EMD-4605 (consensus refinement).

## ACKNOWLEDGMENTS

The work was supported by grants from the MICINN BFU2017-90030-P (to JAH), BFU2017-87316 (to RFL) and by grants from programs Forny (248889/O30) and FRIMEDBIO (261826) from the Research Council of Norway (to AM), the Western Norway Regional Health Authorities (HV projects 911959 to MIF and 912246 to AM) and the K.G. Jebsen foundation (to AM). We thank Kay Diederichs and Pavel Afonine for discussions and help in data processing and refinement, the staff from ALBA and ESRF synchrotron facilities for support during data collection, the Midlands Regional Cryo-EM Facility at LIBSCB, and Christos Savva for assistance during data acquisition. We thank the EM units from CNIO and CNB-CIB (CSIC) for support with the EM facilities and Peter Gimeson from Malvern Panalytical for help with DSC-experiments and discussions. We are very grateful to Prof. Torgeir Flatmark for critical discussions on the manuscript.

