## Supplemental data for "The structure of full-length human phenylalanine hydroxylase in complex with tetrahydrobiopterin"

**This PDF file includes:**

Supplementary text
References for SI reference citations
Figs. S1 to S16
Tables S1 to S3
Captions for movies S1 to S4

**Other supplementary materials for this manuscript include the following:**

Movies S1 to S4

### Supplementary Information

### SI Materials and Methods

**Vectors for expression of hPAH.** Two different constructs of full-length wild-type (WT) human PAH fused N-terminally to maltose-binding protein (MBP) were used in this work. The MBP-(pep)$_{Xa}$-hPAH expressed from a pMal vector (1) is released from the MBP-tag by the protease factor Xa, resulting in a 13 aa truncation at the N-terminus (ΔN13-hPAH) (2). To avoid the secondary cut from factor Xa, the pMAL/hPAH vector was modified to include a tobacco etch virus (TEV)-protease restriction site (ENLYFQ/S) ending with Ser2 of human PAH to avoid extra residues after cutting. The Q5 Site-Directed Mutagenesis Kit (New England Biolabs) was used according to the manual using the forward primer 5'-TATTTTCAGTCTACTGCGGTCCTGGAA-3' and the reverse primer 5'-AAGATTCTCCCTTCCCTCAATCCCGAG-3' (the inserted nucleotides are underlined). MBP-(pep)$_{Xa}$-hPAH-ΔN102/ΔC24 was expressed from vector prepared in (3) and MBP-(pep)$_{Xa}$-hPAH-Y138F in (4). The S23A- and Y377F-hPAH mutations were prepared in the pMAL/hPAH vector using the QuikChange II Site-Directed Mutagenesis kit (Agilent) with appropriate primers.

**Expression and purification of hPAH.** The expression of all PAH proteins was done at 28°C for 16-18 h with 1 mM IPTG. Purification was done in batch mode where the clarified crude extracts were incubated with amylose resin (New England Biolabs) for 3 h and bound fusion protein was eluted with 10 mM maltose. Complete cutting with restriction enzymes required a protease:PAH ratio of 1:300 and 3 h with factor Xa (New England Biolabs) and 1:50 ratio and 2 h for TEV protease. MBP-$(pep)_{Xa}$-ΔN102/ΔC24-hPAH (3) (used to produce the catalytic domain (CD) of hPAH (hPAH-CD)) required 16 h of incubation with Factor Xa. Tetrameric (and dimeric in the case hPAH-CD) PAH proteins were isolated by size exclusion chromatography on a Superdex HiLoad 16/600 200 column (GE Healthcare) in 20 mM Hepes pH 7, 200 mM NaCl. For crystallization, the full-length hPAH proteins were further purified by anion exchange on a HiTrap Q column (GE Healthcare) in 30 mM Tris-HCl pH 7.4, 15% glycerol with a KCl gradient from 30 to 400 mM KCl (5). All purification steps were performed at 4°C. Protein concentration was determined by absorbance at 280 nm using the absorption coefficient $A_{280}$ (1 $mg \cdot ml^{-1} cm^{-1}$) = 1.63 for MBP-hPAH and 1.0 for isolated hPAH.

**Crystallization of hPAH and hPAH-CD.** For full-length hPAH, with regulatory, catalytic and oligomerization domains (RD, CD and OD, respectively) first crystallization screenings were performed by high-throughput techniques in a NanoDrop robot and Innovadyne SD-2 microplates (Innovadyne Technologies Inc.), screening PACT Suite and JCSG Suite (Qiagen), JBScreen Classic 1-4 and 6 (Jena Bioscience) and Crystal Screen, Crystal Screen 2 and Index HT (Hampton Research). Positive conditions were optimized by sitting-hanging-drop vapour diffusion method and the best crystals were obtained in a crystallization condition containing 1.5 M DL Malic Acid pH=7.0, 100 mM Bis-Tris Propane pH=6.2, 0.9 mM Thesit®, 0.98 mM $BH_4$, 25 mM DTT and 1 mM reduced glutathione. PAH was incubated for 5 minutes with an 8-fold molar excess of $BH_4$ in DTT and deposited as 1 µl drops in the crystallization plate wells. Thesit® and reduced glutathione were added directly to the drop before mixing with 1 µL of precipitant solution. The drop was equilibrated against 150 µL of precipitant solution. Protein concentration was assayed at the concentration range of 6-10 mg/mL by mixing 1 µL of protein solution and 1 µL of precipitant solution, equilibrated against 150 µL of precipitant solution. The hPAH-CD at 10 mg/ml in 20 mM Hepes pH 7, 200 mM NaCl was crystallized by the sitting-drop vapor diffusion technique. Crystals were produced by mixing 1 µl of the protein solution and an equal volume of mother liquor (40 mM PIPES, 20% PEG 2000, pH 6.8) as described (6).

**Data collection and structural determination of hPAH and hPAH-CD.** Prior to data collection, crystals were cryoprotected in the precipitant solution supplemented with 25-30% (v/v) glycerol. Diffraction data sets were collected in beamline XALOC at the ALBA synchrotron (CELLS-ALBA, Spain), using a Pilatus 6M detector and a wavelength of 0.97950 Å. Hundreds of crystals were collected for both full-length hPAH and ΔN13-hPAH always presenting very low resolution and strong anisotropy, although ΔN13-hPAH diffracted better than full-length hPAH. Incubation with DTT and $BH_4$ improved diffraction limit up to 2.9 Å resolution along some of the axes. Very likely, the combination of the small truncation of the disordered N-terminus and the high concentration of DTT needed to co-crystallize with $BH_4$ in an aerobic environment were the essential factors that finally allowed the 3D structure of hPAH with all domains to be solved. As previously reported (7), DTT releases iron from the active site of PAH and decreases heterogeneity caused by the redox-active Fe cation. Crystals belong to the C2 monoclinic space group, with unit cell parameters $a$ = 101.94 Å, $b$ = 101.36 Å, $c$ = 203.54 Å, $\alpha = \gamma$ =90º, $\beta$ = 90.00º. Collected datasets were processed with XDS (8) and Aimless (9). Four hPAH monomers were found in the asymmetric unit, yielding a Matthews coefficient of 2.48 $Å^3$/Da (10) and a solvent content of 50.35%. Maximum resolution for refining the model was selected considering both CC1/2 values over 0.30 along the three crystallographic axes and quality of the electron density map after refinement. With these criteria, resolution was chosen at 3.18 Å. The structure was solved by the molecular replacement method using the structure of the catalytic domain of hPAH (PDB ID code 2PAH) as initial model with Morda (a part of ccp4online web services). Refinement of the catalytic domain and manual model building were performed with Phenix (11) and Coot (12), providing electron density map for the regulatory domain (RD) that was included using the equivalent domain in the rPAH (PDB ID code 5DEN). Further refinement was carried out turning on secondary structure restrains for protein and model building that provided a nice electron density map for the complete model (Fig. S1) from residue 22 to 450 except for active-site loop (residues 136-143) that was disordered. Occupancy for the three $BH_4$ molecules was refined yielding values of 0.78 ($BH_4$-1 in chain B), 0.93 ($BH_4$-2 in chain C) and 0.70 ($BH_4$-3 in chain D). In chain D electron density for the region involved in $BH_4$ interaction (residues 247-249 and Phe 254) was compatible with a double conformation with and without $BH_4$. Better refinement statistics were obtained with 0.70 occupancy for the conformation with $BH_4$, and 0.3 occupancy for conformation without $BH_4$. Data collection and processing statistics are shown in Table S1. Crystals for hPAH-CD diffracted up to 1.67 Å resolution and belonged to the C 2 2 $2_1$ space group with unit cell parameters $a$ = 65.85 Å, $b$ = 107.54 Å, $c$ = 124.01 Å, $\alpha = \beta = \gamma = 90^{\circ}$. One monomer of hPAH-CD was found in the asymmetric unit yielding a Matthews coefficient of 3.05 Å$^3$/Da (10) and a solvent content of 59.71%. The structure of the hPAH-CD was solved by molecular replacement using PDB ID 4ANP (13) as initial template. Refinement and manual model building were performed with Phenix (11) and Coot (12), respectively. The stereochemistry of the final models was checked by MolProbity (14). Data collection and processing statistics are shown in Table S1. The atomic coordinates of hPAH and hPAH-CD have been deposited in the Protein Data Bank under codes 6HYC and 6HPO, respectively.

**Cryo-EM sample preparation and data acquisition**. hPAH was thawed to room temperature, diluted to 8 μM in SEC buffer (Hepes 20 mM pH 7, NaCl 200 mM), and submitted to size exclusion chromatography in a 2.3/300 Superdex 200 Increase column (GE) at 4 ºC. Next, 3 μL of the peak fraction were pipetted onto pre-glow-discharged Ultra-Foil Gold 0.6-1 300-mesh grids (Quantifoil), and the excess of liquid was removed using a Vitrobot Mark-III operated at 95% humidity and 4ºC. After blotting, grids were plunged into liquid ethane for vitrification and stored in liquid nitrogen for later use.

**Cryo-EM data acquisition.** Initial screening and optimisation of conditions was performed using an in-house Tecnai-spirit microscope (Thermo Fisher) operated at 120 kV with a TVIPS TemCam-F416 CMOS detector (CNIO EM core facility) and a Talos Arctica (Thermo Fisher) operated at 200 kV equipped with a Falcon III direct electron detector (Centro Nacional de Biotecnología - CSIC). Once optimal conditions were found, samples were imaged at the Midlands Regional cryo-EM facility (Leicester University) on a Titan Krios G3 equipped with a Gatan GIF-quantum energy filter and a Gatan K2 direct electron detector (Thermo Fisher) operated at 300 kV and using a GIF slit of 20 eV. 1494 80-frame uncorrected micrograph movies were acquired using the software EPU (Thermo Fisher) applying image-shift to collect 3 non-overlapping images per foil hole at a magnification of x130,000, yielding a calibrated pixel size of 1.09 Å.px$^{-1}$. Total electron dose per image was 42 e$^-$/Å$^2$ (0.525 e$^-$/Å$^2$.frame).

**Cryo-EM data processing and analysis**. Data processing and analysis was performed using RELION 3.0 (15). Images were imported onto the software package *on-the-fly* and submitted to motion correction using RELION's own implementation of MotionCorr2 (15). During this step, beam induced movement and residual stage drift were corrected for the 80 movie-frame images, using 25 patches (5x5) and applying the detector gain reference image collected before data acquisition. Dose weighting was performed at this stage to compensate for radiation-damage during imaging using a per-frame electron dose of 0.525 e$^-$/Å$^2$. CTF parameters were estimated using Gctf (16). An initial round of reference-free picking using RELION reference-free algorithm and 2D classification was performed to obtain 2D references from a subset of images. Next, 2D references were used for reference-based picking using the whole dataset. The initial set of particles was submitted to several runs of 2D classification to remove false positives from the picking stage, leaving a dataset of 214,017 particles that was further curated using 2D classification to leave a final set of 160,787 high-quality particles (see Fig. S7b). Initial model was generated *ab-initio* using RELION 3.0. We performed initial rounds of 3D refinement and 3D classification to assess the dataset. Due to the high flexibility of the tetramer we decided to process the data using no symmetry (C1). 3D classification using different number of classes yielded several similarly-populated hPAH models resembling the crystallographic tetramer, but featuring different relative positioning of the dimers. A consensus refinement using all 160,787 particles yielded a map with a nominal resolution of 5.3 Å using gold standard Fourier Shell Correlation and a threshold of FSC=0.143 (17) , and a local resolution ranging from 5 to 7 Å. The tetramer could be docked into density showing that the overall structure of the apo-hPAH resembled the crystallographic structure. Nevertheless, the map showed clear signs of heterogeneity for one of the dimers due to flexibility. In order to analyse the flexibility of the monomer we performed a 3D refinement applying a mask to one of the monomers (dotted-lines in Fig. 2C). Using this approach, we obtained a map with improved density for the dimer inside the mask at the expense of the other dimer (see Fig. S7c). Next, we used RELION 3.0 multibody refinement to independently refine the orientations of the particles for the two dimers (dotted-lines in Fig. 2b). This yielded two maps, one for each dimer, with much improved density for the catalytic domains and more homogeneous spread of resolution as seen in Fig. 2b. To further understand the motions of the tetramer, the relative angular assignment of all particles obtained for both dimers during multibody refinement were analysed using a principal component analysis approach with relion_flex_analyse (18). This also allowed us to generate 10 maps for each of the main components describing the relative movement of the dimers within the hPAH tetramer as described in Nakane 2018 (18). B factor sharpening was performed using automatic procedures in RELION. Local resolution was estimated using RELION 3.0. Structures were visualized using UCSF Chimera (19), ChimeraX (20) and Pymol (21). Movies describing the flexibility of hPAH were generated using Pymol (21).

**PAH activity assays.** PAH activity of WT-hPAH and mutants was measured using a Tecan Spark 20M plate reader, with hPAH protein (5 ng/µl) in 20 mM Hepes pH 7, 0.04 mg/ml catalase, 0.5% BSA, 10 µM $(NH_4)Fe(SO)_4$, 1 mM L-Phe, 75 µM $BH_4$ (and 5 mM DTT) at 37 °C in the wells of a Greiner UV-Star® half-area microplate. Three different assays were performed, with three different hPAH states at the start of the reaction: (i) *L-Phe-activated,* hPAH was preincubated with 1 mM L-Phe for 5 min in 100 mM Hepes pH 7, and the reaction was initiated with 75 µM $BH_4$ (with 5 mM DTT) added by the plate reader injector, or (ii) *non-activated* (no ligand during preincubation), hPAH was not preincubated with substrate or cofactor and 1 mM L-Phe and 75 µM $BH_4$ (5 mM DTT) were added simultaneously to start the reaction, or (iii) *pre-catalytic BH4-bound,* preincubated with 75 µM $BH_4$ and 5 mM DTT for 3 min, and L-Phe added to start the assay. The injections were followed by orbital shaking (5 s), and increase in L-Tyr fluorescence was measured with an $\lambda_{ex}$= 274 nm (bandwidth 5.0), $\lambda_{em}$=304 nm (bandwidth 10.0) for 60 s. The standard curves for L-Tyr were prepared with 75 µM $BH_4$ and 5 mM DTT, equilibrated for 5 min 37 ºC, to correct for the inner filter effect of $BH_4$ (22). The initial rate of the reaction was determined by fitting the data points - up to 10 s - to a linear function. PAH specific activity (nmol L-Tyr produced/min/mg PAH at 1 mM L-Phe and 75 µM $BH_4$) was measured with the plate reader assay and corroborated by the HPLC-method as described (23). The steady state kinetic parameters for hPAH, wild-type (WT) and mutants were measured as described (4).

**Differential scanning calorimetry (DSC).** A PEAQ-DSC Automated (Malvern Panalytical) was used to obtain the melting profile of hPAH. In all experiments 19 µM of hPAH in 20 mM Hepes pH 7, 200 mM NaCl was used, with the same buffer as reference, heating from 25-75°C at a scan rate of 200 °C/h. At the scan rate used the aggregation kinetic influence is assumed to be quenched for the duration of the transitions and thus the system is amenable to deconvolution. As the oxidation of $BH_4$ during the DSC scans largely perturbed the base line, 0.5 mM $BH_2$ used and was added to both sample and buffer when indicated. The DSC thermograms were analyzed by PEAQ-DSC analysis software.

**Residue contact analysis.** Residue contacts were calculated for all subunit structures of hPAH using the Bio3D package (24, 25). A pair of residues are considered in contact if the minimal distance between their heavy atoms is ≤4.0 Å. A residue contact is reported on the holo hPAH state only where it is present in at least 2/3 of the structures. For the apo state there was only one structure (1 independent subunit in Tet1 for apo and 1 subunit in Tet1 and 2 in Tet2 for the holo). Differential atomic interactions between apo and holo hPAH are defined as contacts present in one state, but absent in the other state.

**Molecular dynamics simulations.** A total of 16 all-atom unbiased molecular dynamics (MD) simulations were carried out of hPAH and rPAH. Simulations were performed with and without bound $BH_4$, designated $BH_4$ and $BH_4$-free, respectively. The atomic tetramer models were prepared from PDB Tet2 and PDB ID 5DEN, for hPAH and rPAH, respectively, along with the structure of the catalytic domain of hPAH for loop 135-140 (PDB 1J8U) (26). To enhance sampling and statistics each state was simulated four times differing in their generated random initial velocities. All atomic models were prepared with Amber 14 (27) and the corresponding Amber14SB forcefield (28). Parameters for $BH_4$ as well as the catalytic triad and $Fe^{2+}$ were prepared with Antechamber (29) and the general Amber forcefield (GAFF) (30) using a semi empirical model. Protonation states of side chains in the protein and protein-peptide complex were assigned based on the 3D-structure using the well-established software PROPKA at pH 7.0 (31). For each of the simulations, the system was neutralized using a mixture of $Cl^-$ and $Na^+$ counter ions, and the protein was solvated in a periodic truncated octahedron box with TIP3 water molecules (32), providing 16 Å of water between the protein surface and the periodic box edge. The solute was minimized for 5 000 steps, followed by 5 000 steps of minimization of the whole system with restraints on the tetramerization helix, and finally 5 000 steps minimization of all atoms. The protein was then heated to 100 K with weak restraints for 20 ps, and to 300 K for 1 ns. Equilibration with constant pressure and temperature (NPT) of the system was performed for a total of 1.5 ns prior to the production with reduced restraints on the solute. The production runs lasted for 100 ns and were performed with (1) two simulations with constant volume and energy (NVE; *ntt=0*), and (2) two simulations with constant temperature (*Berendsen* thermostat for temperature control (*ntt=1*)) and constant volume (NPE). All simulations were run with a 1 fs time step, using SHAKE constraints on hydrogen-heavy atom bonds and increased SHAKE and EWALD tolerance (*tol = 0.000001*). All simulations were run with GPU acceleration (33) on Tesla K20Xm cards. The simulations were analyzed using ptraj (34) and Bio3D (24, 25). Fluctuation profiles are shown at monomer level, averaged over the 4 subunits from 4 simulations. Wilcoxon signed-rank test were performed to test for significant differences in fluctuations between apo and holo / hPAH and rPAH simulations. Targeted MD (TMD) simulations were performed using PDB Tet2 (with $BH_4$ and L-Phe) as starting structure, and the substrate bound conformation of the catalytic domain (PDB ID 1MMK (35) as target structure. All-atomic tetrameric models for TMD were prepared using Bio3D and Amber 14. The starting structure was equilibrated using the same protocol as described above. The TMD simulation was performed in Amber 14 using a 1 fs time step and Langevin thermostat (*ntt=3*) with constant temperature and volume (NVP) over a period of 2 ns. All heavy atoms of the CD of chain A (residues 125-407) were used for calculating the RMSD, and hence the restraint forces, driving the conformational change from an RMSD of 2.85 to 0.5 Å between the starting and target structures.

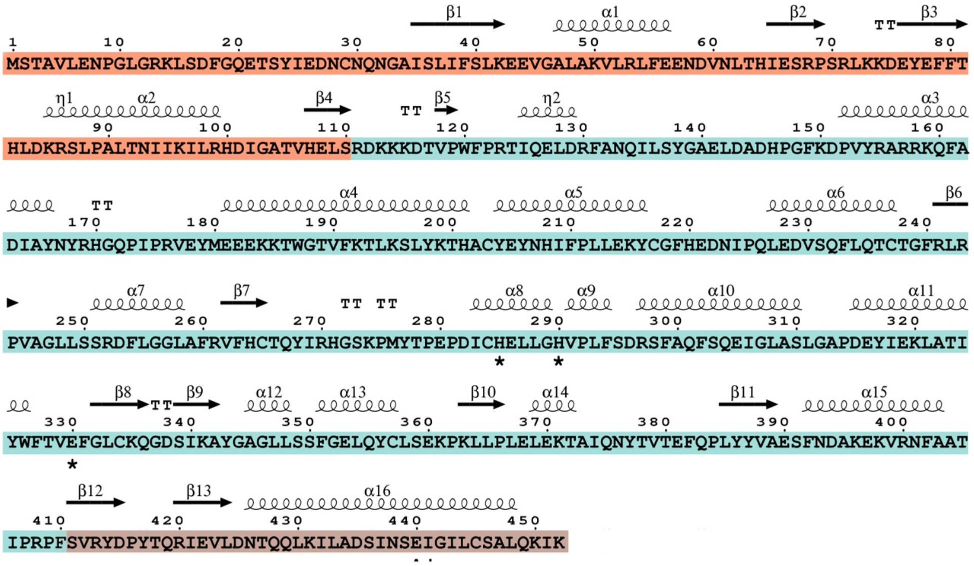

**Fig. S1. Sequence for hPAH.** Primary sequence of hPAH in which the secondary structure elements are indicated and numbered. Asterisks indicate residues coordinating catalytic Fe ion. Domains are colored differently (RD in orange, CD in cyan and OD in brown).

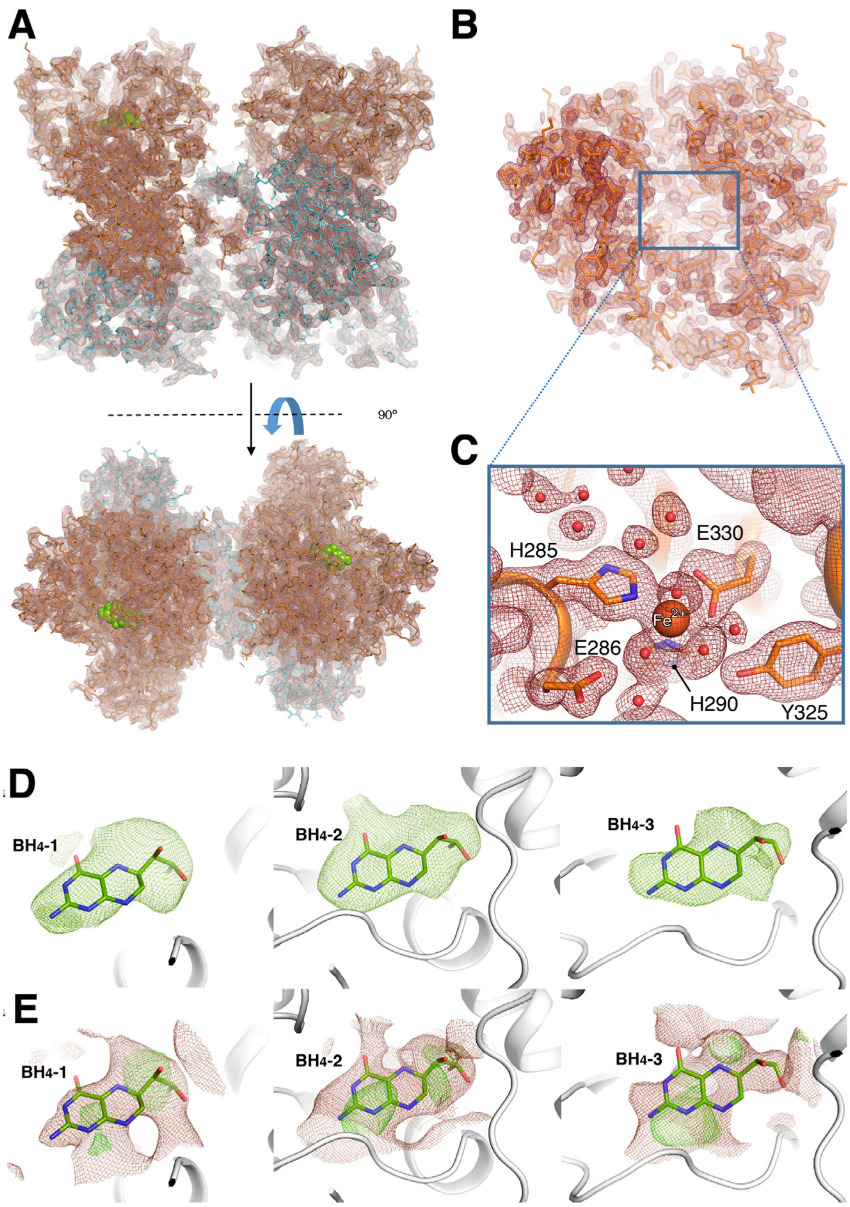


**Fig. S2. Electron density map for hPAH.** (**A**) Electron density map for full-length hPAH (2Fo-Fc map contoured at 1σ) in complex with $BH_4$. The structure corresponds to Tet1, which presents the $BH_4$ (green spheres) bound in two monomers (depicted in orange sticks) and the other two without the cofactor (depicted in cyan stick). (**B**) Electron density map for hPAH-CD (2Fo-Fc map contoured at 1σ) in complex with iron. (**C**) Detailed view of the boxed area shown in (B). Residues found in close proximity to iron are labeled and depicted as capped sticks**.** Water molecules are shown as red spheres. (**D**) Polder OMIT maps (in green) for each $BH_4$ (depicted as capped sticks) calculated by setting the solvent exclusion radio to 3Å and contoured at 4σ. (**E**) 2Fo-Fc maps (coloured in brown and contoured at 0.8σ) and initial Fo-Fc maps (coloured in green and contoured at 2.5σ) for each $BH_4$ molecule. $BH_4$-1 is the cofactor found in chain B of the Tet1 whereas $BH_4$-2 and $BH_4$-3 are found in chains C and D of the Tet2. Occupancies determined after refinement for each $BH_4$ molecule were 0.78 ($BH_4$-1), 0.93 ($BH_4$-2) and 0.70 ($BH_4$-3).

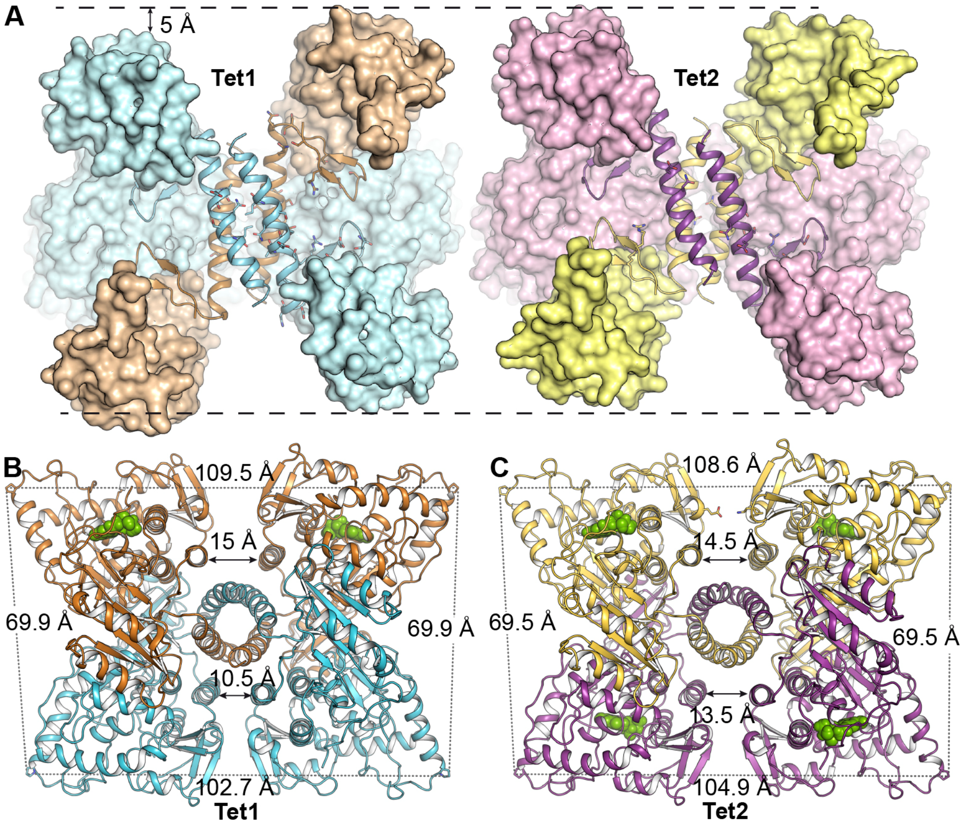


**Fig. S3. Oligomeric arrangement in the apo-holo (Tet1) and holo-holo (Tet2) tetramers of hPAH**. (**A**) Different $BH_4$ occupancy results in gliding of the dimers inside the tetramer through the oligomerization helices. Each hPAH monomer is represented by its molecular surface except for the OD depicted as ribbon. Two monomers in the tetramer have been partially omitted for clarity reasons. Ribbon representation of Tet1 (**B**) and Tet2 (**C**). Tetramers are encased in an isosceles trapezoid (109.5 Å and 102.7 Å for apo-holo tetramer and 108.6 Å and 104.9 Å for holo-holo tetramer, distances measured between residue P152). Distances between equivalent monomers in the tetramer also change depending on $BH_4$ occupancy. Some representative distances, between Cα of K398 from a monomer and D394 from the symmetry-related monomer, are indicated by a double arrow. Color code for chains in Tet1 and Tet2 are conserved among the panels. It is worth to mention a significant shortening of the distance between apo monomers (chains A, B in Tet1) *vs*. holo monomers (chains C, D in Tet1 and chains A, B and C, D in Tet2) (Fig. 2B, 2C), thus showing that in the absence of cofactor equivalent monomers are closer to each other and that they separate upon $BH_4$ binding.

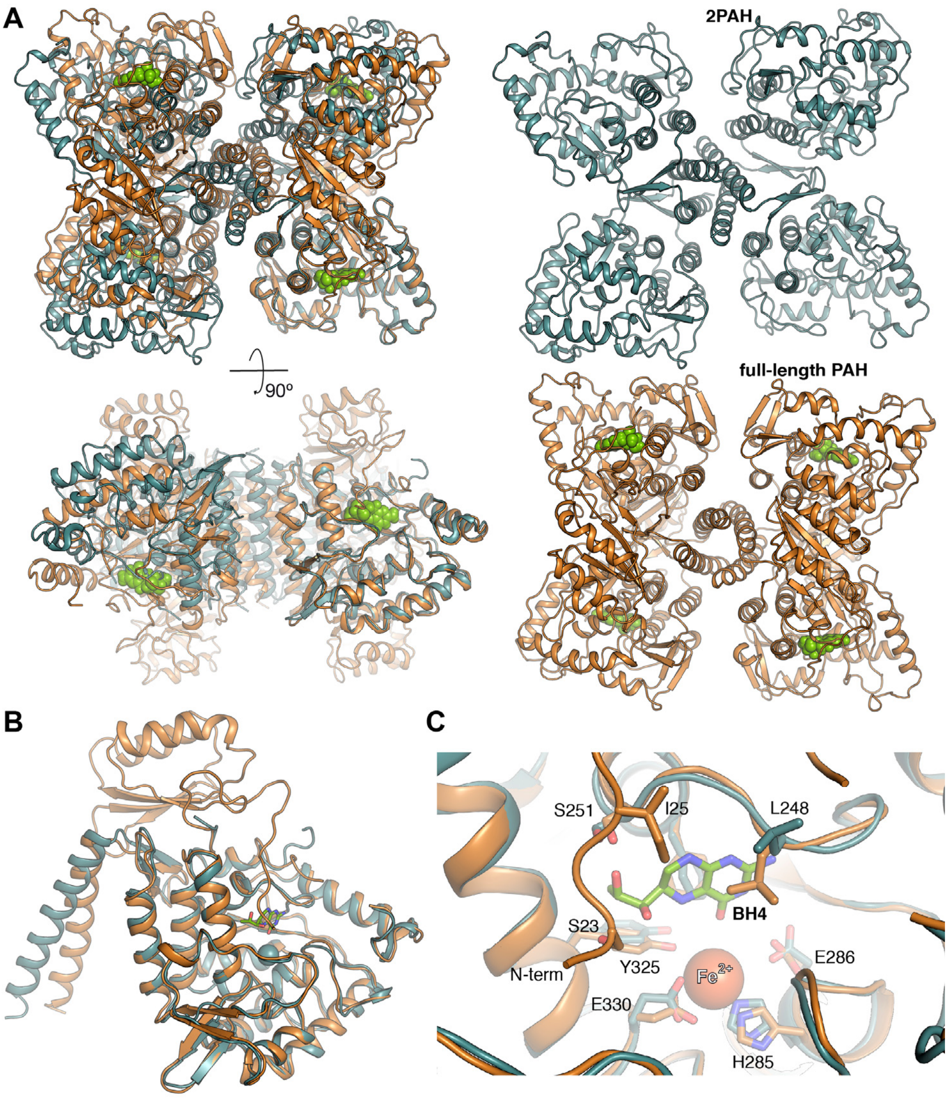


**Fig. S4. Structural comparison of full-length human PAH and the truncated version lacking RD.** (**A**) Structural superimposition of full-length hPAH (Tet1 colored ochre, this work) and apo hPAH without RD (residues 118-452; blue, PDB ID code 2PAH). The truncated structure showed an asymmetric arrangement of the dimers. (**B**) Structural superimposition of full-length hPAH (this work) and truncated 2PAH monomers. While the CD is very similar (RMSD 1.13 Å) to that in truncated PAH, the RD imposes a different angle on the OD helix. (**C**) Detailed view of the catalytic site in both structures. Relevant residues are drawn as capped sticks and labeled.

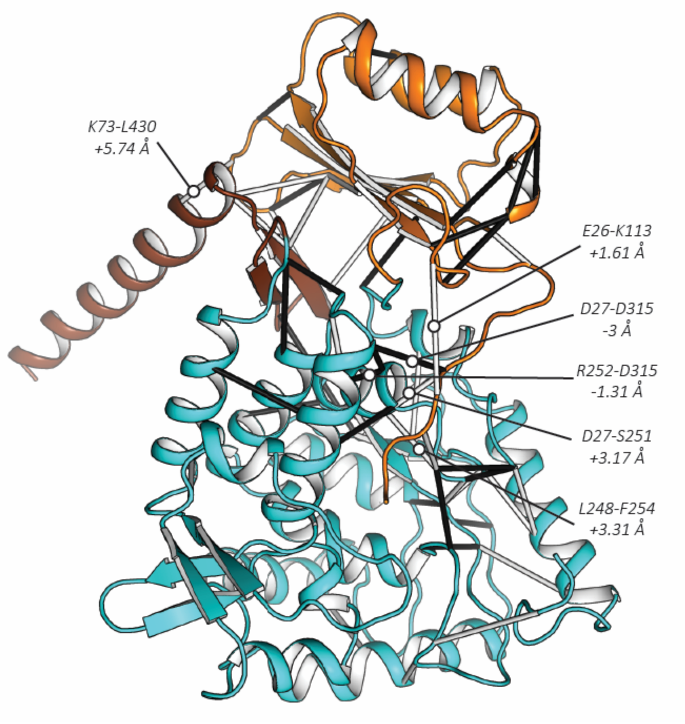


**Fig. S5. Differential atomic interactions in hPAH monomer upon $BH_4$-binding**. The six largest differential atomic interactions, defined as residues being in contact (≤ 4 Å) in one state but not in the other (see Table S2), are depicted with solid sticks between residues. White and black sticks depict contacts in apo-hPAH and holo-hPAH, respectively. All-atom distances between all pairs of residues are calculated with the available monomer for apo-hPAH (equivalent C/D chains in Tet1) while the distances in holo-hPAH are the average of three distances in the three monomers available (chains A/B in Tet1, chains A/B in Tet2 and chains C/D in Tet2). Distance differences mentioned in the main text are labelled on the structure with the change in Å from apo-hPAH to holo-hPAH. Distances involving residues 137-142 are not included as they are missing in the structures.

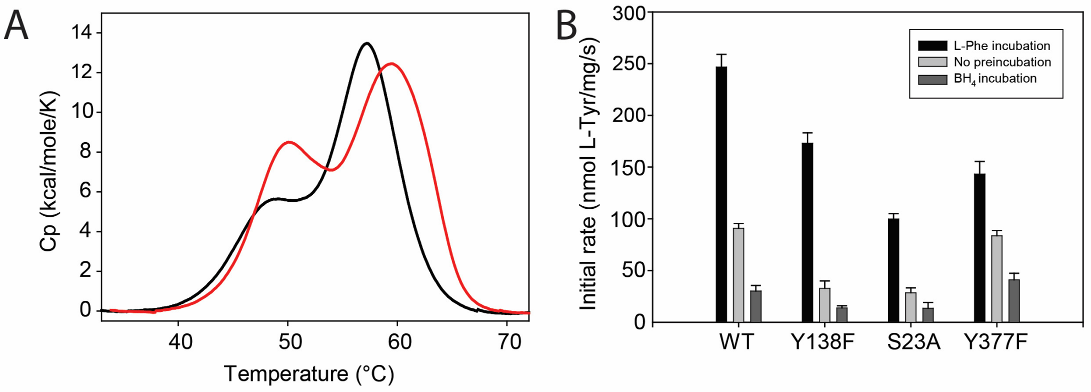


| $T_m$ (°C) | $\Delta H_{cal}$ (kcal/mol) | $\Delta H_{VH}$ (kcal/mol) | $\Delta H_{VH}/\Delta H_{cal}$ |
|---|---|---|---|
| no ligand | | | |
| 48.3 ± 0.2 | 60.9 ± 4.1 | 87.4 ± 6.0 | 1.4 |
| 57.8 ± 0.1 | 95.4 ± 1.9 | 113.7 ± 0.9 | 1.2 |
| + $BH_2$ | | | |
| 50.1 ± 0.1 | 69.0 ± 2.2 | 93.8 ± 3.8 | 1.4 |
| 59.3 ± 0.1 | 101.0 ± 2.1 | 110.0 ± 0.8 | 1.1 |

C

| hPAH | $V_{max}$ (nmol L-Tyr/min/mg) | $K_m(BH_4)$ (µM) | $S_{0.5}$(L-Phe), ($h$) (µM) |
|---|---|---|---|
| WT | 4745 ± 245 | 35 ± 5 | 170 ± 10, (2) |
| Y138F | 3621± 232 | 35 ± 5 | 172 ± 7, (1.7) |
| S23A | 2098 ± 110 | 27 ± 1 | 680 ± 59, (1.3) |
| Y377F | 3447 ± 256 | 53 ± 4 | 704 ± 48, (2) |

**Fig. S6. Thermal stability of apo- and holo-hPAH and initial reaction rates and steady state catalytic parameters of WT and hPAH mutants. (A)** Differential scanning calorimetry (DSC) thermograms of hPAH (19 µM subunit) without (black) and with (red) 0.5 mM 7,8-dihydrobiopterin ($BH_2$), added as an oxidized, non-reactive $BH_4$ analog. The curves are representative from 3 scans, heating from 25 to 70 °C at a scan rate of 200 °C/h. The table shows midpoint melting temperature ($T_m$), calorimetric enthalpy ($\Delta H_{cal}$), Van't Hoff enthalpy ($\Delta H_{VH}$) and $\Delta H_{VH}/\Delta H_{cal}$ ratio derived from the deconvolution analysis of the DSC thermograms for the thermal denaturation of hPAH in the apo and $BH_2$-bound states applying a non-2-state model. Values are average and SD from triplicate measurements. Standard error from fittings were <0.1% for $T_m$-values and <1% for ΔH-values. (**B**) Initial PAH reaction rates for WT-hPAH, and the Y138F, S23A and Y377F mutants, at three preincubation conditions (see Materials and Methods and Fig. 3 in main text). The initial rates are calculated by fitting the activity (nmol L-Tyr/mg/s) at 37 ºC from 0-10 s to a linear function. (**C**) Steady-state kinetic parameters for the hPAH WT and mutants; the substrate concentrations were 1 mM L-Phe ($BH_4$ variable) and 75 µM $BH_4$ (L-Phe variable, up to 1 mM L-Phe), determined at 25 °C with 5 min preincubation with the corresponding concentration of L-Phe. $S_{0.5}$(L-Phe) represents the L-Phe concentration at half-maximal activity and $h$ is the Hill coefficient obtained from the sigmoidal dependence of the activity on L-Phe concentration. $K_m(BH_4)$ and $S_{0.5}$(L-Phe) for Y138F have been previously reported (4).

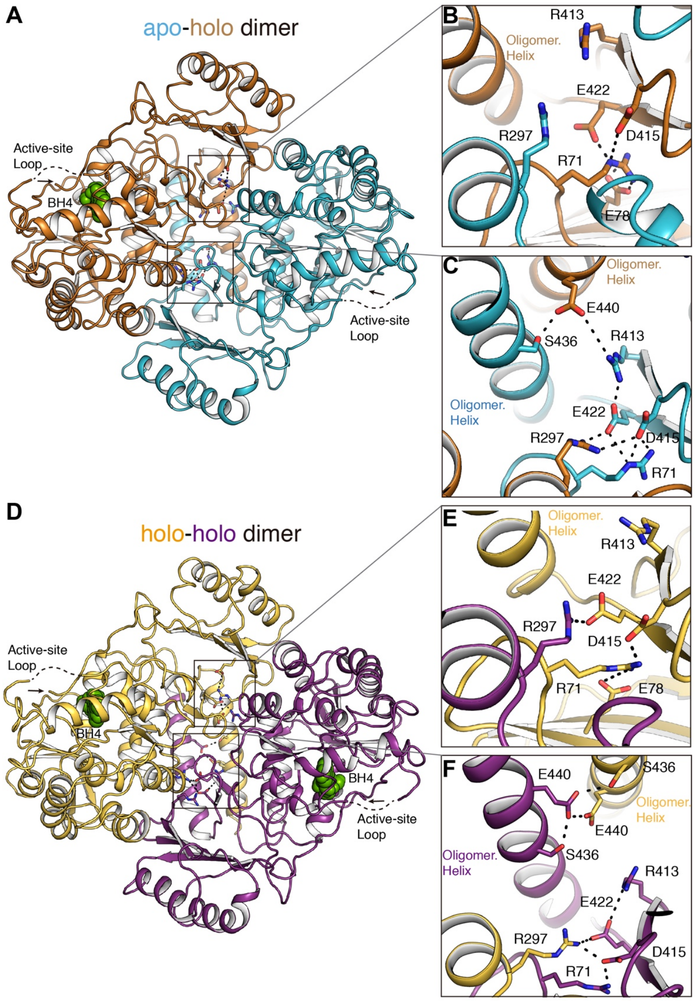


**Fig. S7. Intra dimer interactions in apo-holo (Tet1) and holo-holo dimers (Tet2) of hPAH**. (**A**) Ribbon representation of the apo-holo dimer of Tet1. Monomers are colored differently (apo in cyan and holo in orange) with relevant residues depicted as capped sticks and $BH_4$ as green spheres. Active site loop is labeled and N-terminus indicated by an arrow. (**B**,**C**) Detailed view of the apo-holo dimer interface showing the network of salt-bridge interactions. While the R297 (CD) residue of the holo hPAH establishes a strong network of salt-bridge interactions with the E422, D415 (OD) and R71 (RD) of the partner (panel C), the same R297 residue in the apo hPAH does not participate in such network (panel B). Very interestingly, the network observed in the apo hPAH connects the core residues (E422 and D415) with the OD helices of both chains in the dimer (panel C). Thus, the salt bridge network connects residues R297(holo) - E422(apo) - R413(apo) - E440(holo); finally, E440, in the OD helix, makes a polar interaction with S436(apo), also in the OD helix of the partner. (**D**) Ribbon representation of the holo-holo dimer of Tet2. Active site loop is labeled and N-terminus indicated by an arrow. (**E**,**F**) Detailed view of the holo-holo dimer interface showing the network of salt-bridge interactions. In the case of holo-holo dimer there is no asymmetry in the intermolecular interactions and the R297 residue, in both chains, makes salt-bridge interactions with the acidic residues E422 and D415 of the partner (panels E, F). Moreover, contrary to the apo-holo dimer, there is no interaction between the CD and the OD helices. Remarkably, both helices in the dimer are connected by polar interactions through the S436 and E440 residues of the two chains (panel F).

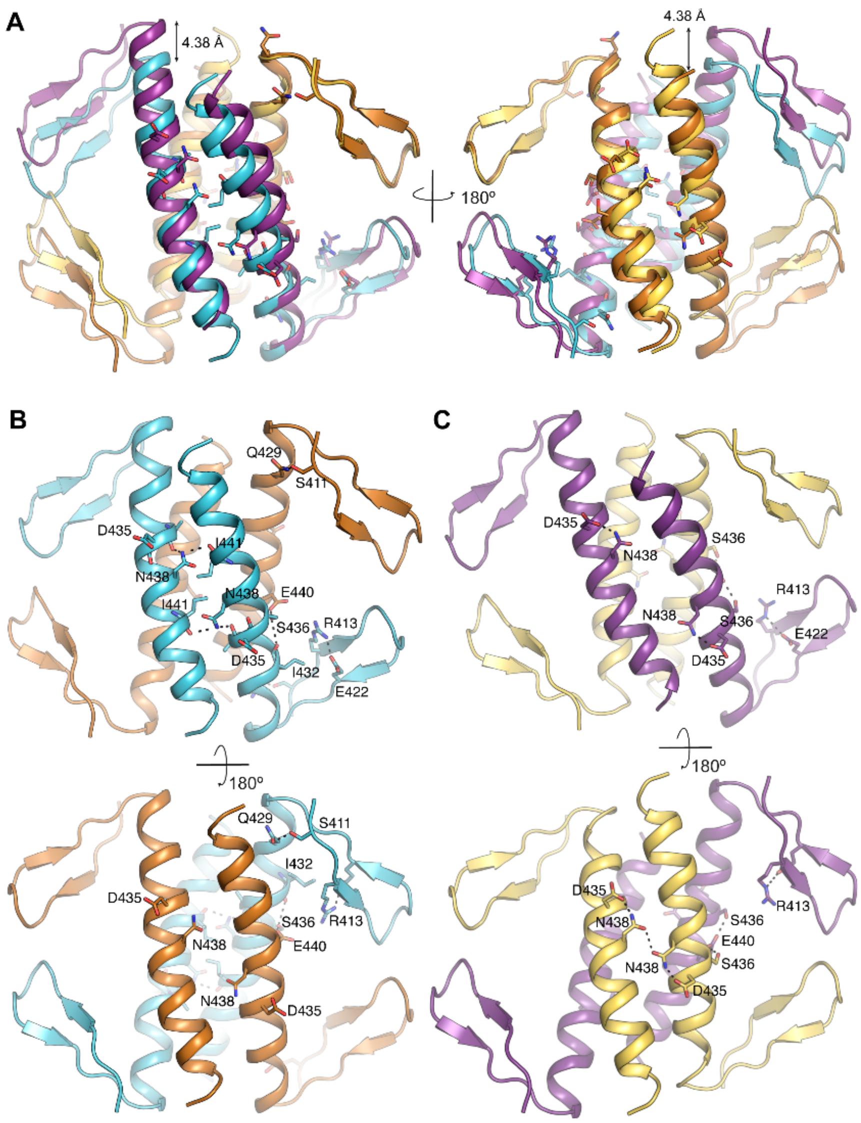


**Fig. S8. Intermolecular interactions in oligomerization domains of Tet1 and Tet2 of hPAH.** Color code as in Figure 2 of main text. (**A**) Structural superposition of ODs of Tet1 and Tet2. Relevant residues (as detailed in panels B and C) depicted as capped sticks. Gliding along the OD helices is indicated by a double arrow (see main text for details). (**B**) Intermolecular polar interactions (dashed lines) in the OD of Tet1. (**C**) Intermolecular polar interactions (dashed lines) in the OD of Tet2.

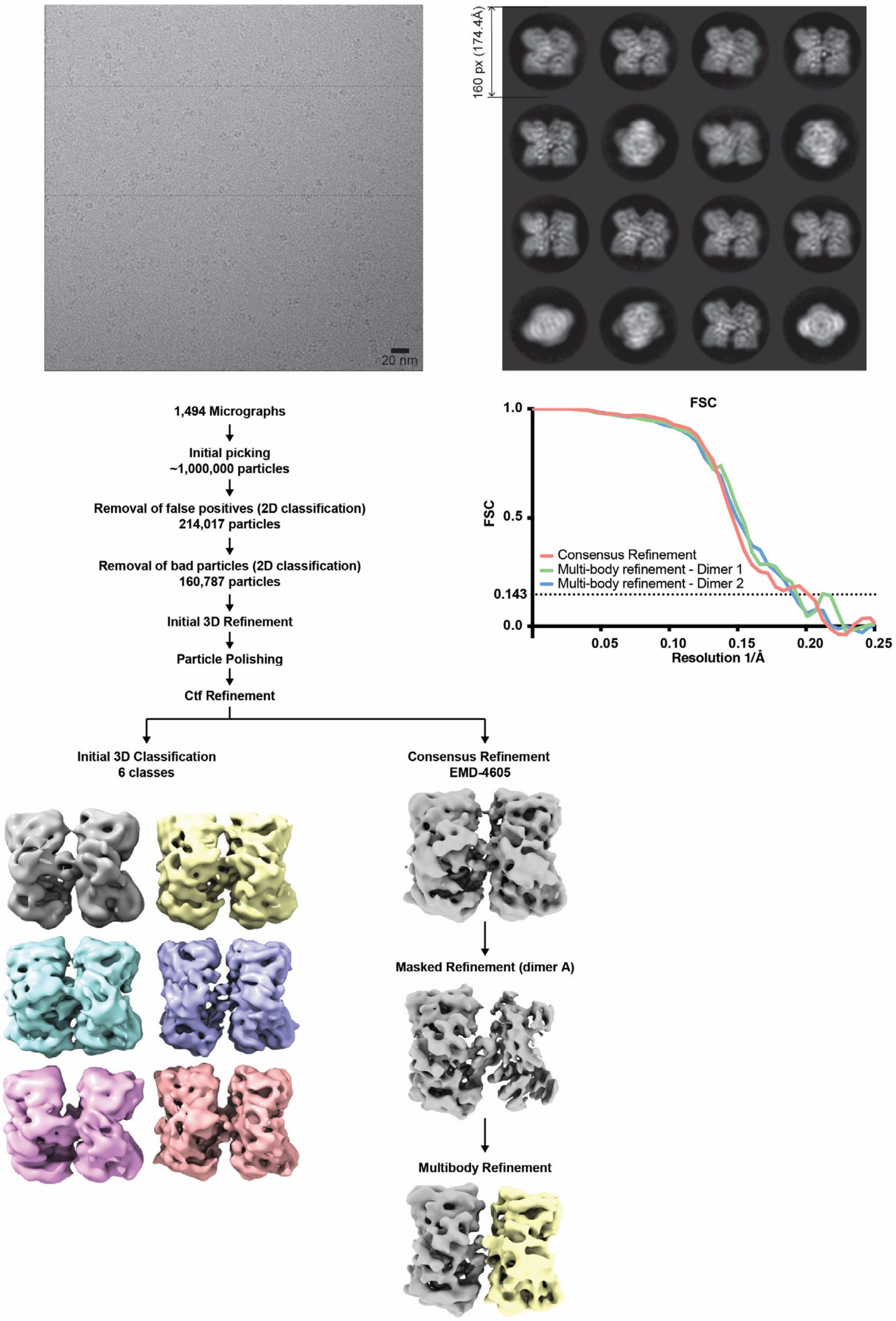


**Fig. S9. hPAH cryo-EM data analysis. (A)** Representative micrograph from the hPAH cryo-EM dataset. **(B)** Class averages resulting from 2D classification of the curated selection of 160,787 particles. **(C)** Data processing overview. After initial picking and curation of the dataset, particles were submitted to 3D classification (left) and to refinement followed by multibody refinement and flexibility analysis (see Materials & Methods for details). **(D)** Fourier Shell Correlation (FSC) for the consensus refinement and multibody-refinement maps. Lines represent correlation between two independent halves of the dataset for each map (gold-standard FSC). The horizontal dashed-line shows the resolution cut-off (0.143).

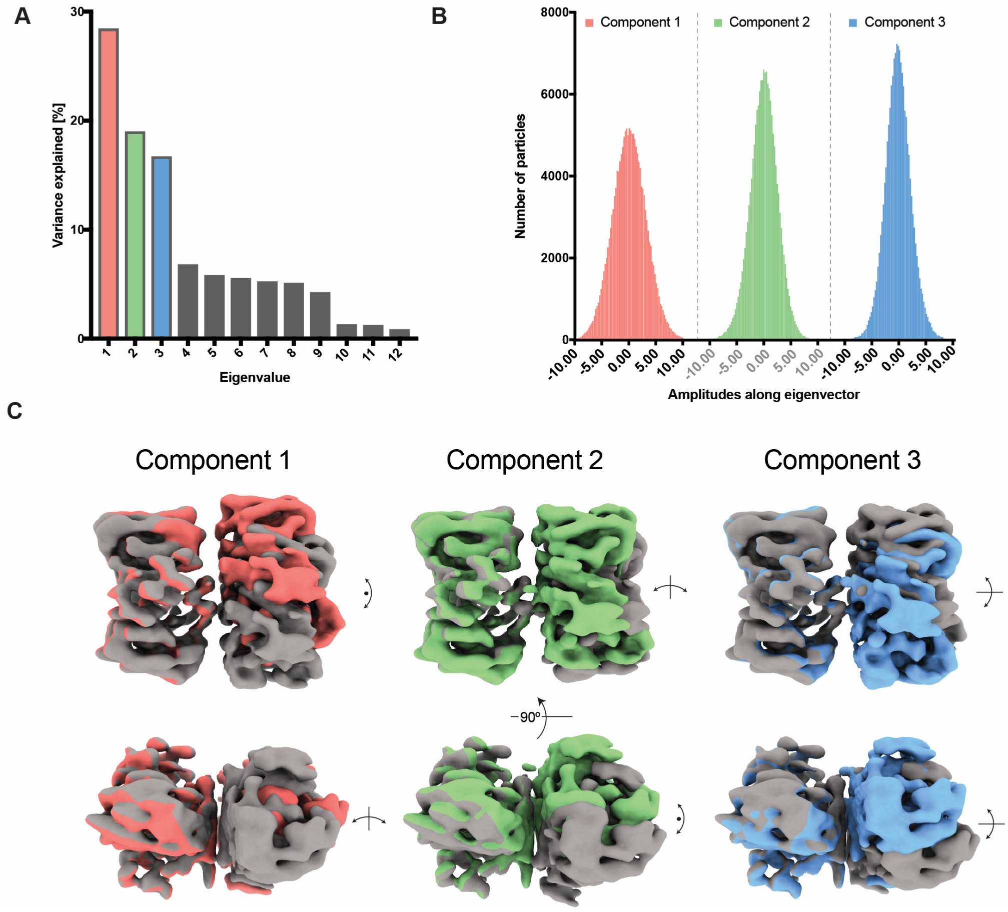


**Fig. S10. hPAH flexibility analysis. (A)** Results from the Principal Component Analysis of the relative orientations calculated for the two dimers during multi-body refinement. Bars represent the amount of variance within the dataset explained by each of the components. The three first eigenvectors explain more than 60% of the variance observed and are highlighted in color. **(B)** Distribution of amplitudes for all particles along the three eigenvectors highlighted in A. Note that distributions are monomodal, suggesting a continuous movement of the dimers. **(C)** After PCA flexibility analysis in RELION, particles were pooled into 10 equally populated bins for each of the components. For each bin, a map was generated representing the median orientation of the particles within that group. Panel B shows a superimposition of the first and tenth map for each of the three main components, highlighting the flexibility range of apo-hPAH. See SI Material & Methods for details on the flexibility analysis.

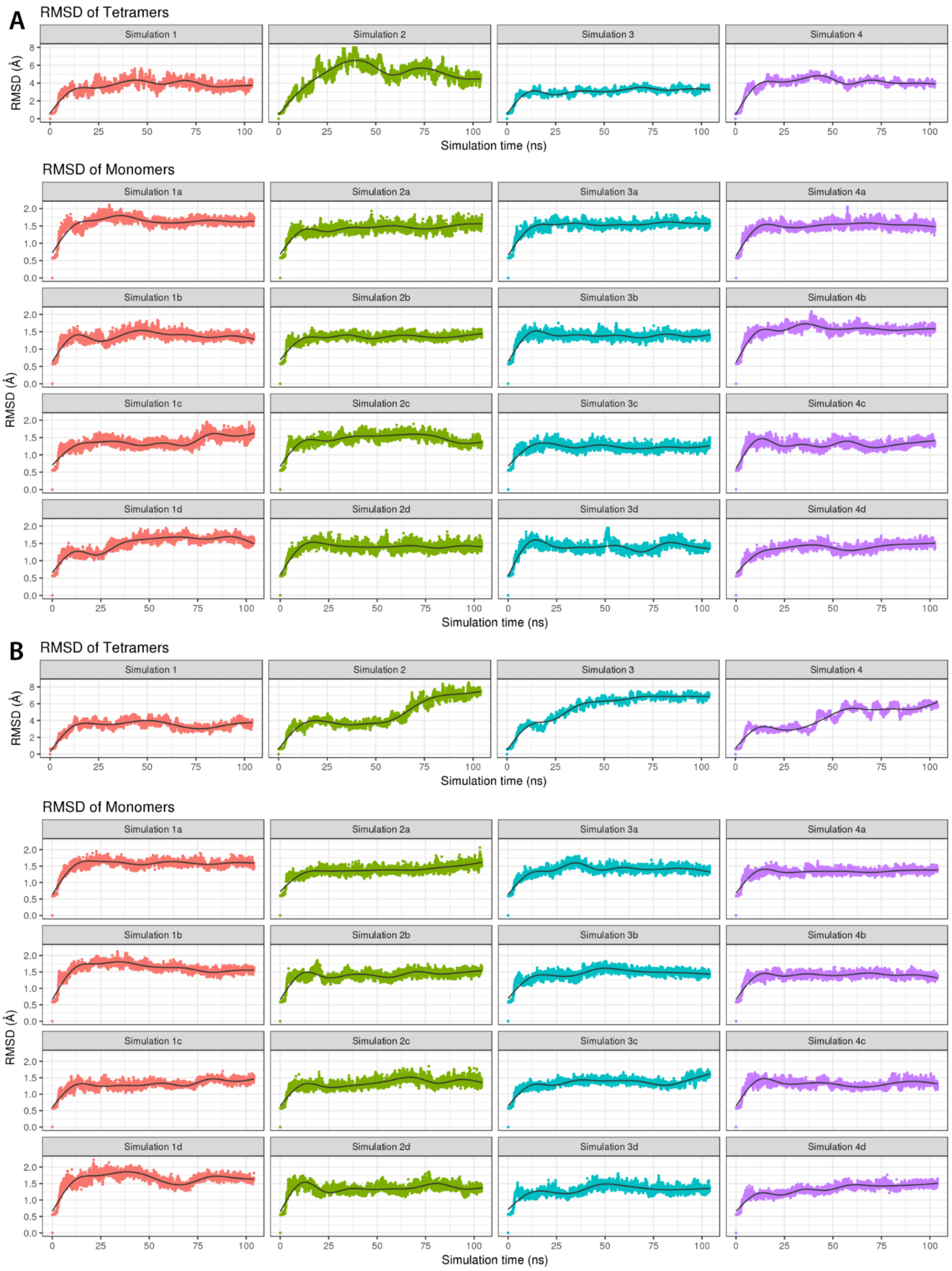


**Fig. S11. Structural deviations along MD simulations of hPAH**. The figure shows the root mean square deviation (RMSD) towards the structure of hPAH (Tet2) along the 8 100 ns-long MD simulations. The RMSD is calculated for the entire tetramer (upper panel) and for the individual subunits (lower panel). RMSD values are shown for (**A**) apo and (**B**) $BH_4$-bound.

A

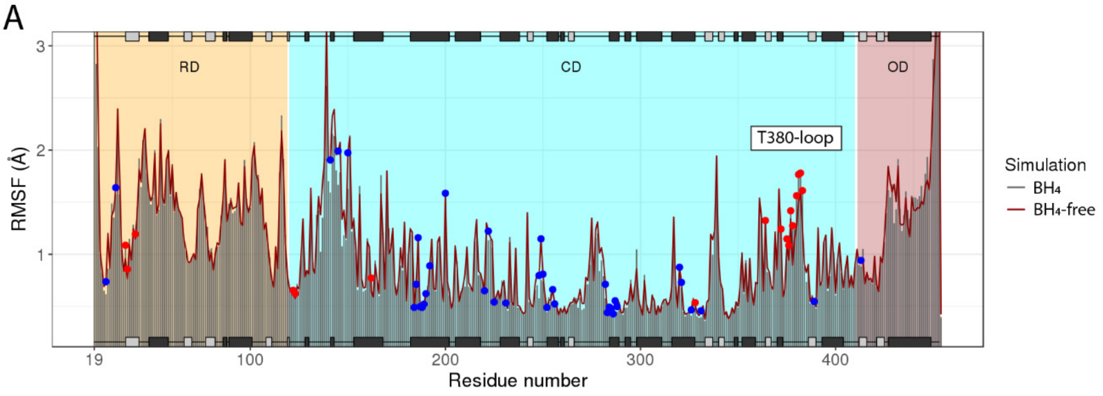


B

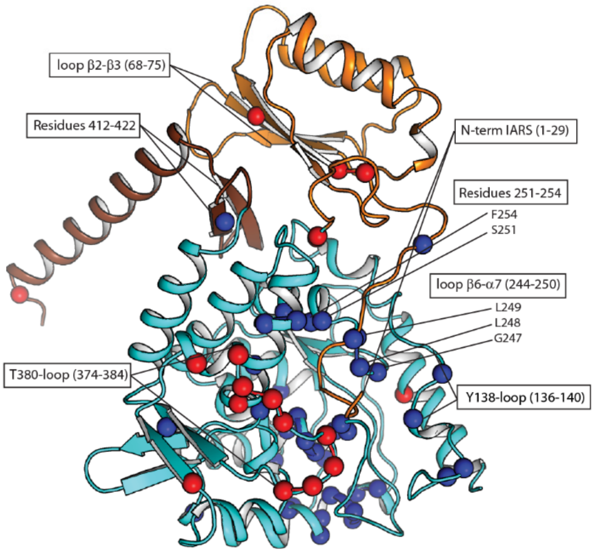


**Fig. S12. Analysis of residue fluctuations measured by MD simulations of hPAH with and without $BH_4$.** (**A**) Fluctuations (root-mean-square; RMSF) of residues in simulations with $BH_4$ (red line) or without $BH_4$ (gray bars). Residues showing significant differences in fluctuation amplitude ($p < 0.05$) upon $BH_4$-binding are marked with dots (blue: smaller fluctuations with $BH_4$, red: larger fluctuations with $BH_4$). (**B**) Residues showing significant differences between apo and holo are visualized on hPAH monomer. Blue and red residues are less and more mobile upon $BH_4$-binding, respectively. The positions of the Y138- and T380-loops and several hotspots for PKU mutations (see main text for details) are indicated. RD, CD, OD stand for regulatory-, catalytic- and oligomerization domains, respectively.

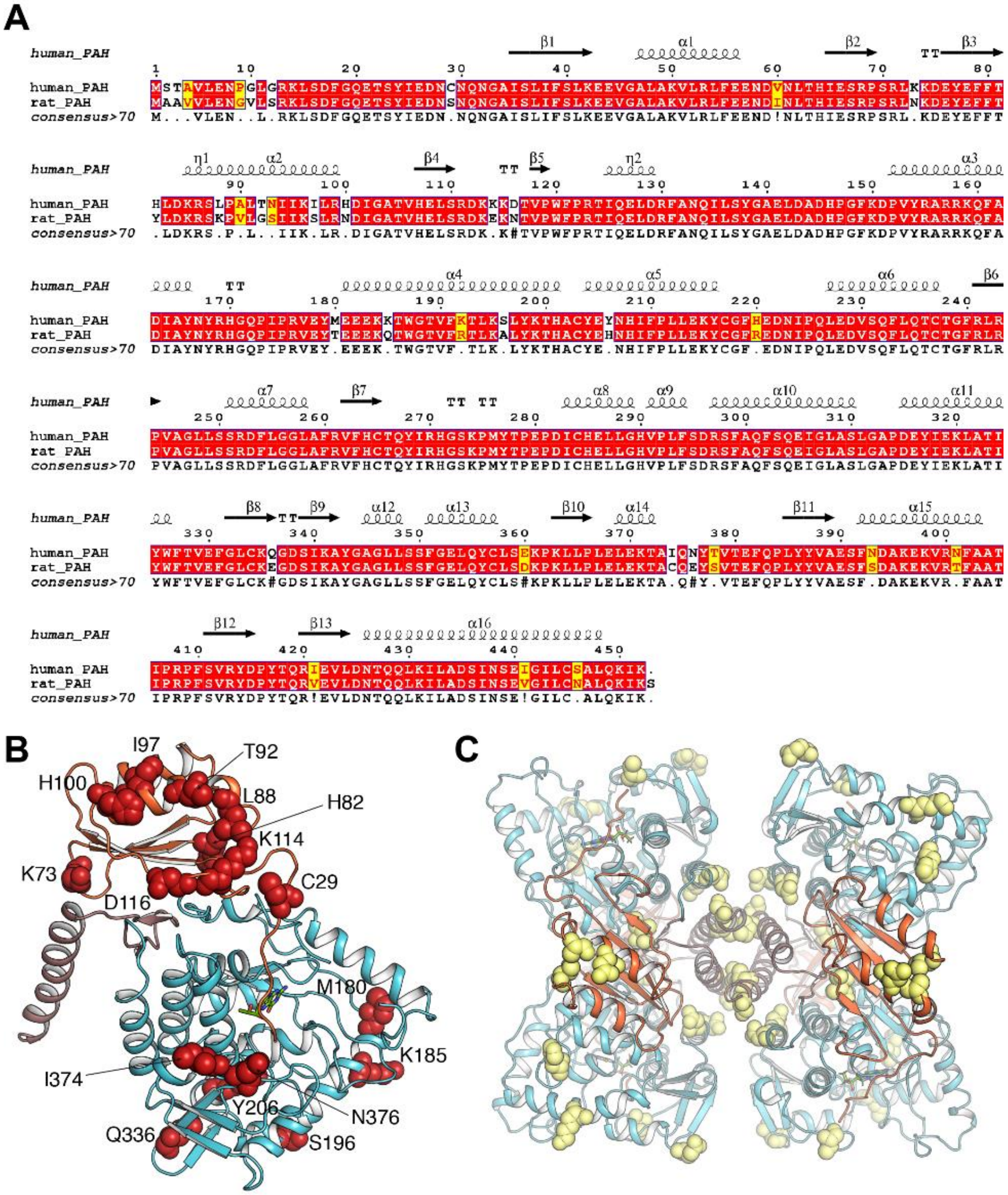


**Fig. S13. Mutations in human PAH *vs.* rat PAH. (A)** Sequence comparison of human (h) and rat (r) PAH. **(B)** Distribution of non-conservative mutations between hPAH and rPAH. Residues in hPAH are depicted as red spheres and labeled. Domains are colored as in Figure 1B. (**C**) Distribution of conservative mutations on the hPAH tetramer. Mutated residues are depicted as yellow spheres. Protein is color-coded as in Figure 1B.

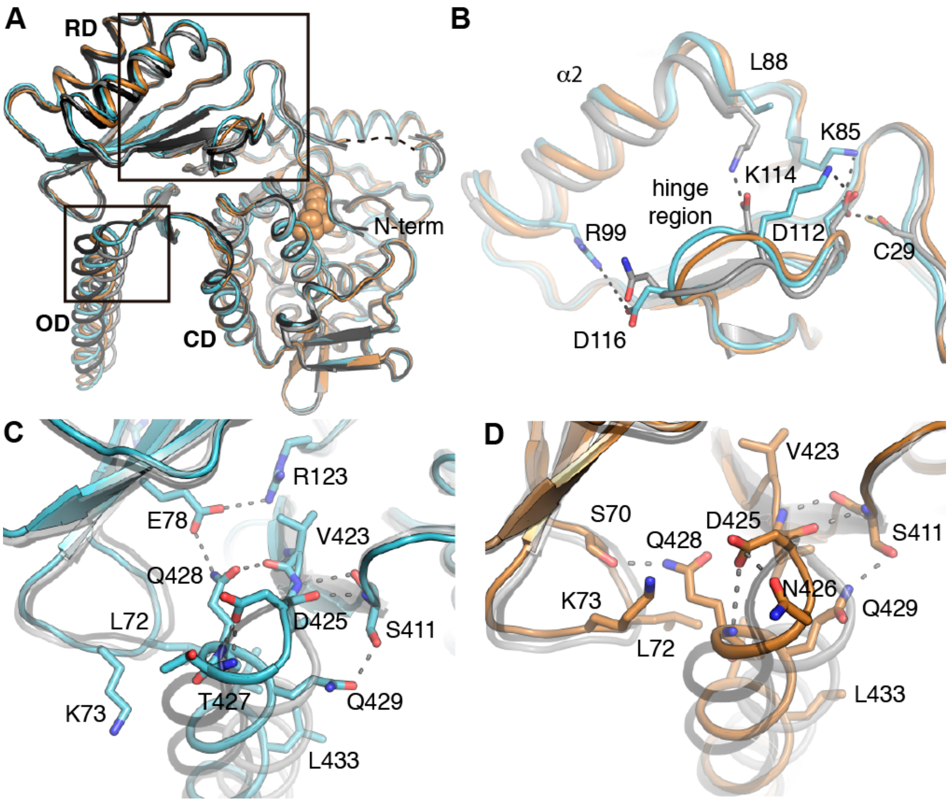


**Fig. S14. Structural differences between human and rat PAH monomers**. (**A**) Structural superimposition of apo hPAH (light blue), holo hPAH (orange) and the two conformers observed in apo rPAH (gray and black). Regions presenting most relevant changes are highlighted in boxes. The $BH_4$ cofactor is represented as spheres. (**B**) Detailed view of the structural changes observed in the N-terminal domain. (**C**) Detailed view of the interactions in the oligomerization helix of the apo hPAH. Relevant residues are drawn as capped sticks and labeled. (**D**) Detailed view of the interactions in the oligomerization helix of the holo hPAH.

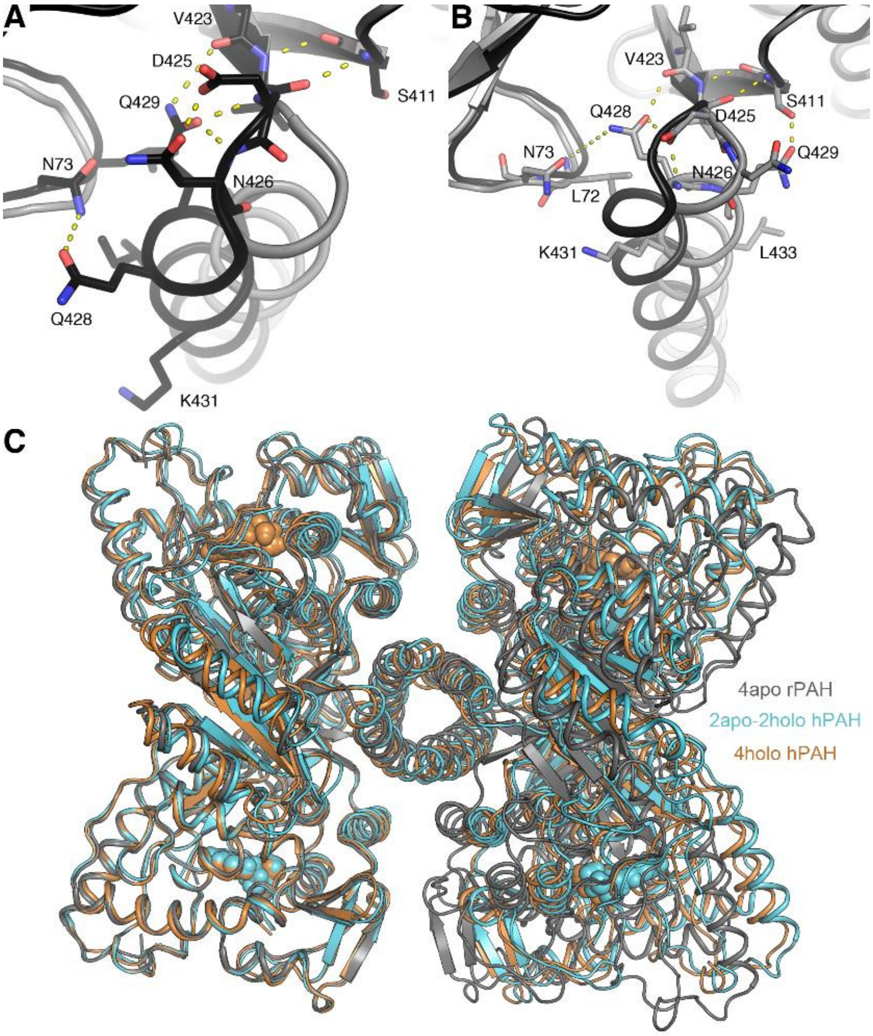


**Fig. S15. Structural comparison of human PAH *versus* rat PAH.** (**A**) Detail of the interactions observed in one conformer of the apo rPAH (black; PDB code 5DEN) in the OD-helix region. Residues involved are represented as capped sticks and labeled. The backbone of the other observed conformer (gray) is shown for comparison. (**B**) Detail of the interactions in the other conformer of the apo rPAH (gray; PDB code 5DEN) in the OD-helix region. Residues involved are represented as capped sticks and labeled. The backbone of the other conformer (black) is show for comparison. (**C**) Structural superposition of the tetrameric arrangement observed in apo rPAH (gray, PDB code 5DEN) and in apo-holo and holo hPAH tetramers (cyan and ochre respectively, this work).

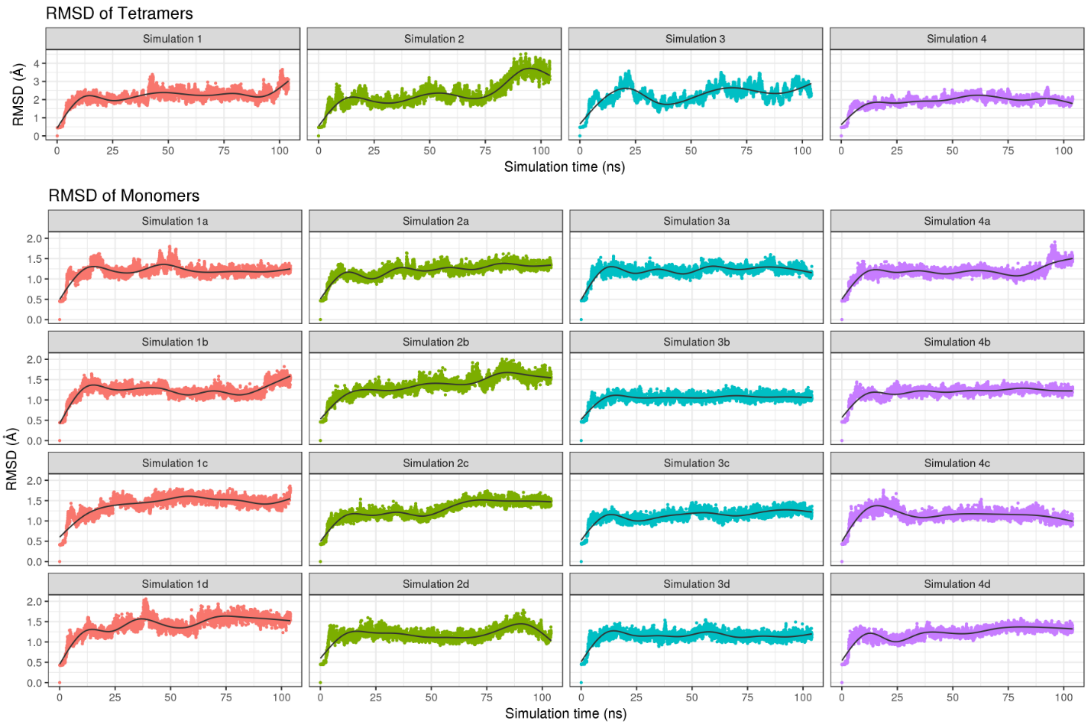


**Fig. S16. Structural deviations along MD simulations of rPAH**. The figure shows the root mean square deviation (RMSD) towards the structure of rPAH (PDB ID 5DEN) along the 4 100 ns-long MD simulations. The RMSD is calculated for the entire tetramer (upper panel) and for the individual subunits (lower panel).

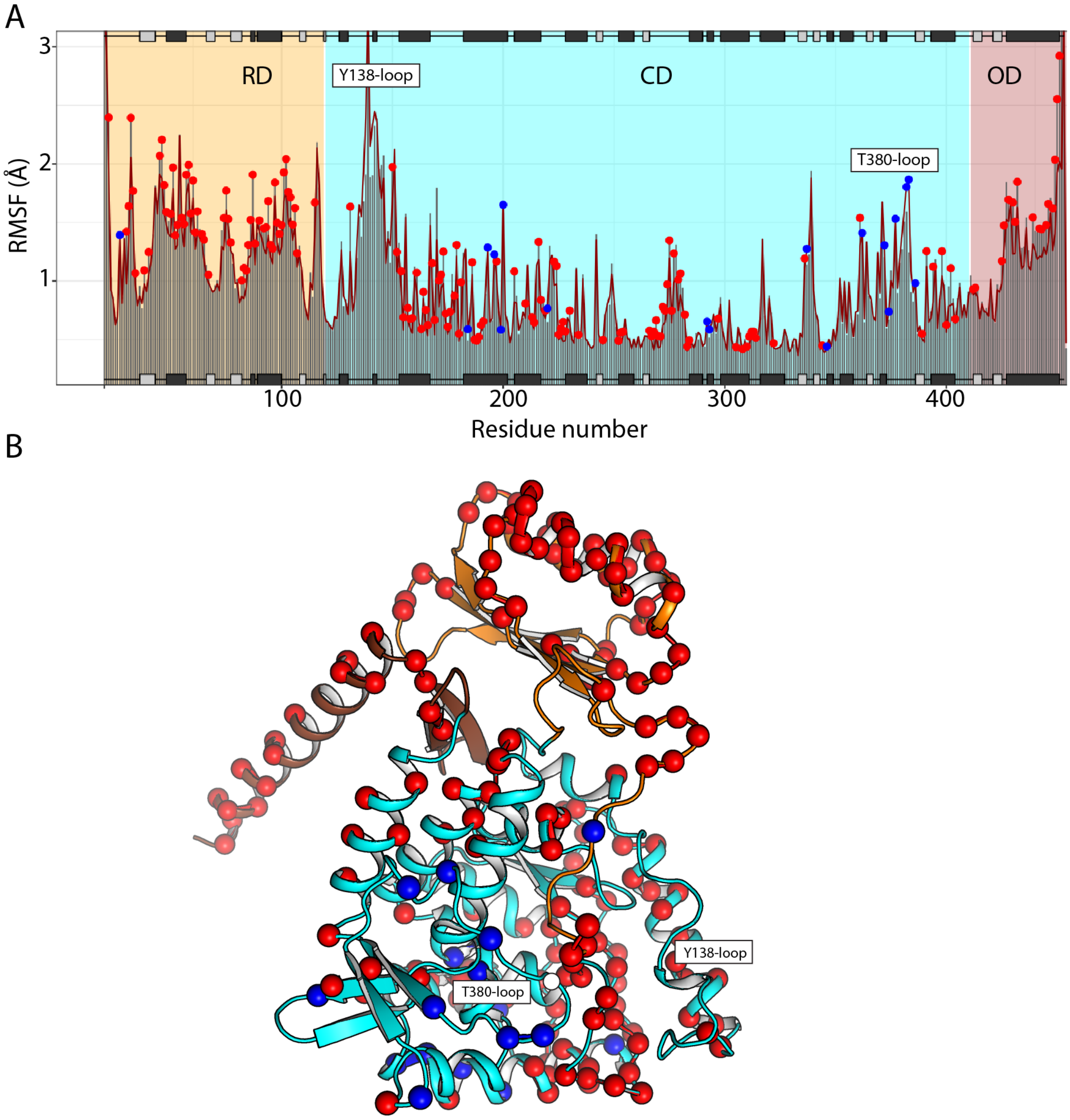


**Fig. S17. Analysis of residue fluctuations measured by MD simulations of hPAH and rPAH.** (**A**) Fluctuations (root-mean-squared fluctuations; RMSF) of residues in simulations with hPAH or rPAH. Residues showing significant differences in fluctuation amplitude ($p < 0.05$) between rPAH and hPAH are marked with dots (blue: smaller fluctuations in hPAH, red: larger fluctuations in hPAH). (**B**) Residues showing significant differences between hPAH and rPAH are visualized on a hPAH monomer. Blue and red residues are those that are less and more mobile in hPAH, respectively. The position of the Y-138 and T380-loops are indicated.

**Table S1. Crystallographic data collection and refinement statistics***

| | hPAH full-length | hPAH-CD |
|---|---|---|
| **Data collection** | | |
| Wavelength (Å) | 0.97926 | 0.97924 |
| Space group | C2 | $C222_1$ |
| Unit cell *a, b, c* (Å) | 101.94, 101.37, 203.54 | 65.85, 107.54, 124.01 |
| Unit cell α, β, γ (º) | 90, 90.00, 90 | 90, 90, 90 |
| T (K) | 100 | 100 |
| X-ray source | Synchrotron | Synchrotron |
| Resolution range (Å) | 45.57–(3.29–3.18) | 32.77–(1.73–1.67) |
| Unique reflections | 34280 (3926) | 586507 (30774) |
| Completeness (%) | 94.3 (83.6) | 100.00 (100.00) |
| Multiplicity | 2.9 (2.5) | 11.4 (11.8) |
| $<I/\sigma(I)>$ | 2.4 (0.30) | 16.2 (1.6) |
| *CC1/2* | 1.00 (0.42) | 1.00 (0.66) |
| **Refinement** | | |
| Resolution range (Å) | 33.92–3.18 | 32.77–1.67 |
| $R_{work}/R_{free}{}^a$ | 0.2619/ 0.3107 | 0.1607/ 0.1770 |
| No. Atoms | | |
| Protein | 13792 | 2889 |
| Water | 5 | 295 |
| Ligand | 51 | 1 |
| **R.m.s. deviations** | | |
| Bond length (Å) | 0.007 | 0.007 |
| Bond angles (°) | 1.25 | 1.15 |
| **Ramachandran** Favored/outliers (%) | 92.55/0.24 | 97.40/0.00 |
| Monomers per AU | 4 | 1 |
| **PDB code** | 6HYC | 6HPO |

*Values between parentheses correspond to the highest resolution shells

$^aR_{work}/R_{free} = \Sigma_{hkl}| F_o - F_c | / \Sigma_{hkl} | F_o |$, where $F_c$ is the calculated and $F_o$ is the observed structure factor amplitude of reflection hkl for the working / free (5%) set, respectively.

**Table S2. Comparative analysis of distances upon $BH_4$-binding.** The distances are measured between the closest atoms from residue 1 and 2 in a subunit. The distances measured for apo-hPAH are from the single monomer available (equivalent C/D chains in Tet1) while the distances in holo-hPAH are the mean of three distances in the three monomers available (chains A/B in Tet1, chains A/B in Tet2 and chains C/D in Tet2).

| | **Residue 1** | **Domain** | **Residue 2** | **Domain** | **Distance in apo-hPAH (Å)** | **Distance in holo-hPAH (Å)** | **Change in distance upon $BH_4$-binding (Å)** |
|---|---|---|---|---|---|---|---|
| **1** | LYS73 | R | LEU430 | O | 3.34 | 9.08 | +5.74 |
| **2** | LEU248 | C | PHE254 | C | 3.57 | 6.88 | +3.31 |
| **3** | LEU321 | C | PHE402 | C | 7.08 | 3.84 | -3.24 |
| **4** | ASP27 | R | SER251 | C | 2.49 | 5.66 | +3.17 |
| **5** | ASP145 | C | TYR277 | C | 3.46 | 6.50 | +3.03 |
| **6** | ASP27 | R | ASP315 | C | 6.32 | 3.32 | -3.00 |
| **7** | LYS85 | R | ARG111 | C | 6.45 | 3.82 | -2.63 |
| **8** | LEU248 | C | GLU280 | C | 6.20 | 3.66 | -2.53 |
| **9** | GLU78 | R | GLN428 | O | 3.69 | 5.76 | +2.07 |
| **10** | LEU136 | C | ARG158 | C | 3.11 | 5.07 | +1.96 |
| **11** | LEU136 | C | GLY247 | C | 5.30 | 3.35 | -1.95 |
| **12** | ARG241 | C | TYR414 | O | 6.38 | 4.44 | -1.94 |
| **13** | VAL177 | C | GLN267 | C | 5.66 | 3.73 | -1.92 |
| **14** | ILE38 | R | GLU78 | R | 3.49 | 5.31 | +1.83 |
| **15** | LEU136 | C | LEU248 | C | 5.17 | 3.35 | -1.82 |
| **16** | ARG71 | R | GLU78 | R | 4.97 | 3.17 | -1.80 |
| **17** | ALA246 | C | THR266 | C | 3.36 | 5.16 | +1.80 |
| **18** | GLY247 | C | THR266 | C | 3.64 | 5.38 | +1.74 |
| **19** | TYR154 | C | TYR268 | C | 3.09 | 4.79 | +1.70 |
| **20** | LYS85 | R | SER110 | R | 5.30 | 3.62 | -1.68 |
| **21** | VAL423 | O | GLN428 | O | 2.62 | 4.27 | +1.65 |
| **22** | GLU26 | R | LYS113 | C | 3.76 | 5.37 | +1.61 |
| **23** | ALA313 | C | ILE318 | C | 3.74 | 5.31 | +1.56 |
| **24** | SER251 | C | ALA322 | C | 5.54 | 4.02 | -1.53 |
| **25** | THR189 | C | MET276 | C | 3.20 | 4.60 | +1.40 |
| **26** | ARG252 | C | ILE318 | C | 4.40 | 3.00 | -1.40 |
| **27** | ARG86 | R | PRO89 | R | 4.36 | 2.98 | -1.38 |
| **28** | ILE35 | R | LEU88 | R | 4.75 | 3.38 | -1.37 |
| **29** | PHE191 | C | GLU214 | C | 3.76 | 5.13 | +1.37 |
| **30** | ARG261 | C | GLN301 | C | 5.09 | 3.73 | -1.36 |
| **31** | ALA309 | C | ARG408 | C | 4.13 | 2.78 | -1.35 |
| **32** | SER36 | R | ARG123 | C | 5.23 | 3.92 | -1.31 |
| **33** | ARG252 | C | ASP315 | C | 4.58 | 3.27 | -1.31 |
| **34** | GLU66 | R | ARG420 | O | 2.24 | 3.54 | +1.30 |
| **35** | LYS85 | R | PRO89 | R | 4.95 | 3.65 | -1.30 |

**Table S3. Cryo-electron microscopy data collection and reconstruction statistics for apo hPAH.**

| Cryo-EM | apo hPAH |
|---|---|
| **Data collection** | |
| Microscope | FEI Titan Krios (Leicester University) |
| Detector | Gatan K2 Summit (counting) |
| Magnification | 130000 |
| Voltage (kV) | 300 |
| Total Electron dose ($e^-$/Å$^2$) | 42 |
| Number of movie frames | 80 |
| Defocus range (µm) | -1 to -4 |
| Pixel Size (Å) | 1.09 |
| **Reconstruction** | |
| Symmetry imposed | C1 |
| Initial micrographs (no.) | 1492 |
| Initial particle images (no.) | 214017 |
| Final particle images (no.) | 160787 |
| Box size (pixel) | 160 |
| FSC threshold | 0.143 |
| Map resolution (Å) | 5 |
| Map resolution range (Å) | 5-7 |
| Map sharpening B factor (Å$^2$) | -313 |
| PDB Accession code | EMD-4605 |

- **Movie S1. Structural changes from Tet1 to Tet2 in human PAH.** Morphing between the tetramer presenting the $BH_4$ bound in half of the active sites (Tet1) and the tetramer with $BH_4$ bound to all four active sites (Tet2). View of the hPAH along the OD helices

- **Movie S2. Structural changes from Tet1 to Tet2 in human PAH.** Morphing between the tetramer presenting the $BH_4$ bound in half of the active sites (Tet1) and the tetramer with $BH_4$ bound to all four active sites (Tet2). Lateral view showing gliding along the OD helices between Tet1 and Tet2.
- **Movie S3. Principal component analysis of tetramer flexibility in apo hPAH.** After PCA flexibility analysis in RELION, particles were pooled into 10 equally populated bins for each of the components. For each bin, a map was generated representing the median orientation of the particles within that group. Movie S1 cycles through the ten maps for each of the three main components, highlighting the flexibility range of apo-hPAH. See SI Material & Methods for details on the flexibility analysis.
- **Movie S4. Targeted Molecular Dynamic (TMD) simulation.** The movie shows the trajectory of the TMD simulation of Tet 2 in complex with L-Phe and $BH_4$ guided towards the target substrate-bound structure defined by the ternary hPAH-CD:$BH_4$:THA complex (PDB code 1MMK). The catalytic domain is shown in cartoon, with Y138, S23 and $BH_4$ highlighted in sticks and balls, and residues 22-26 in orange.
