## Supplementary figures and images for "The structure of full-length human phenylalanine hydroxylase in complex with tetrahydrobiopterin"

### Supplemental movie 3

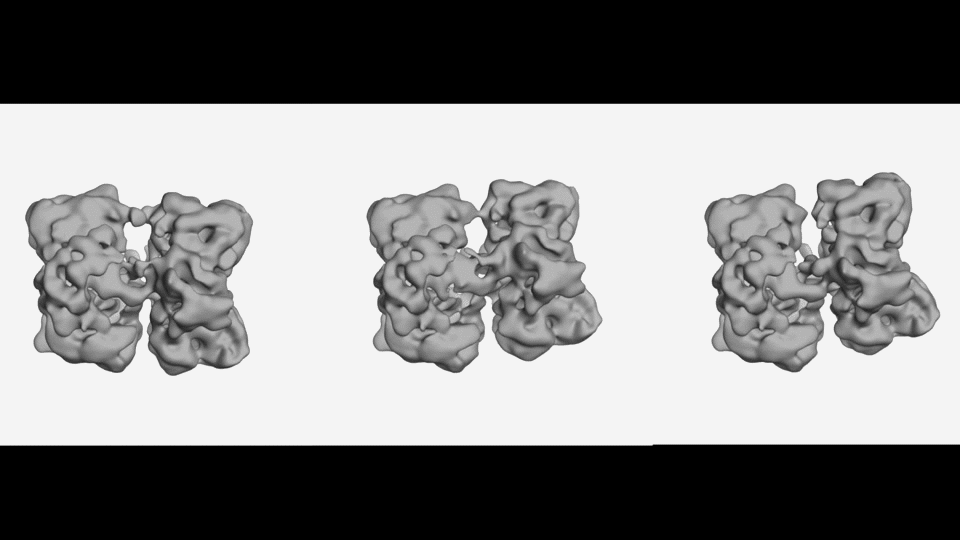
